# Cell type-specific representation of spatial context in the rat prefrontal cortex

**DOI:** 10.1101/2023.10.31.564949

**Authors:** H Brünner, H Kim, S Ährlund-Richter, J van Lunteren, AP Crestani, K Meletis, M Carlén

**Affiliations:** Department of Neuroscience, Karolinska Institutet, Stockholm, Sweden; Department of Neuroscience and Behavioral Sciences, University of São Paulo, Ribeirão Preto, São Paulo, Brazil; Picower Institute of Learning and Memory, Massachusetts Institute of Technology, Cambridge, MA, USA; Scientifica Ltd, London, UK; Center for Neuroscience, University of California, Davis, CA, USA

## Abstract

The ability to represent one’s own position in relation to cues, goals, or threats is crucial to successful goal-directed behavior. Using transgenic rats expressing Cre recombinase in parvalbumin (PV) neurons (PV-Cre rats) we demonstrate cell type-specific encoding of spatial and movement variables in the medial prefrontal cortex (mPFC) during reward foraging. Single neurons encoded the conjunction of the animal’s spatial position and the location of the reward, referred to as the spatial context. The spatial context was most prominently represented by the inhibitory PV interneurons. Movement towards the reward was signified by increased local field potential (LFP) oscillations in the gamma band but this LFP signature was not related to the spatial information in the neuronal firing. The results highlight how spatial information is incorporated into cognitive operations in the mPFC. The presented PV-Cre line opens for expanded research approaches in rats.

## Introduction

Navigating the environment is fundamental to animals’ foraging for sustenance. Foraging decisions incorporate the own localization and place-associated objects, rewards, and threats. The importance of the hippocampal formation in spatial processing has been firmly established, but spatial activation patterns are also widespread in cortical regions^1–5^. This suggests that spatial variables are diversely utilized in brain functioning, although the specificities remain to be established. The mPFC serves as a hub for integrating different information streams subserving cognitive computations^6^. Many cognitive tasks engaging the mPFC require the incorporation of spatial contingencies, and single mPFC neurons have been shown to merge spatial information with other relevant task variables, for example, goals, past choices, or task rules^7–9^. Thus, the context and/or relevance of a location appears to be represented in the mPFC^10^. Cortical information processing involves dynamic interactions between excitatory and inhibitory neurons organized in network motifs (for review see^11,12^). The role of inhibition and the network computations employed by inhibitory interneurons have at large been disregarded in studies of spatial processing in the mPFC^5,7,9^.

Interneurons expressing the calcium-binding protein parvalbumin (PV) constitute the largest inhibitory population both in cortex and in the hippocampus^11,12^. This population is characterized by fast, non-adapting firing and synaptic targeting of perisomatic sites close to the site of action potential (AP) generation in local excitatory neurons. This gives PV interneurons strong influence over the excitatory output, and the ability to coordinate local population activities at both behavioral timescales (seconds) and at oscillation timescales (milliseconds (ms) to hundreds of ms)^13^. Over the past 40 years, a wealth of theoretical and experimental research has provided evidence for the central role of fast-spiking PV interneurons (FS-PV neurons) in gamma oscillations (30-80 Hz)^14^, ubiquitous rhythms that typically emerge in brain regions rich in inhibitory interneurons providing perisomatic inhibition. These transient rhythms appear spontaneously during sleep and waking states but also during distinct processing^15,16^. Hippocampal gamma rhythms are associated with spatial sequences and movement variables^17,18^ and while gamma rhythms in the PFC have been intensively studied in relation to cognitive operations, the involvement in spatial processing is largely unexplored.

Much knowledge on spatial processing, and learning and memory, has been derived from studies in freely moving rats^19–22^, which tolerate more dense and bulky recording technologies than mice. Rats are considered to better reflect human physiology, disease, and behavior than mice^23^, but the utility of this model animal has decreased in response to the rapid expansion of the genetic and experimental toolbox for studies in mice. We here introduce a transgenic knock-in rat line expressing Cre recombinase in PV neurons (PV-Cre) throughout the cerebrum and cerebellum. This tool enables *ex vivo* or *in vivo* identification and manipulation of select PV populations in rats. To characterize the involvement of FS-PV neurons in prefrontal spatial processing we employed chronic electrophysiological recordings and optotagging in adult PV-Cre rats engaging in a self-paced, asymmetric reward-seeking task on a linear track. One end of the track was attached to a reward platform and the other end to a trigger platform void of reward. To trigger reward delivery in the reward platform the animals needed to visit the trigger platform, leading the animals to ambulate across the track. Generalized linear models (GLMs) identified significant encoding of both spatial and movement variables in FS-PV neurons, other narrow-spiking (NS) neurons, and wide-spiking (WS) putative excitatory neurons, respectively. The animal’s spatial position on the track was prominently represented in the neuronal activity of all three populations, with subpopulations of single neurons encoding both the animal’s spatial position on the track and if the animal was moving towards or away from the reward. The context of the track traversal was particularly evident in the activities of the mPFC FS-PV neurons. Furthermore, the traversals to reward versus (vs) to trigger were associated with significant differences in the LFP activity in the gamma band (30-50 Hz). However, we did not find support for local gamma oscillations organizing PrL spatial representation.

## RESULTS

### Specific targeting of PV neurons in PV-Cre rats

To allow study and manipulation of PV neurons in rats, we used CRISPR/Cas technology to generate a knock-in rat line with Cre expression in PV neurons (**Figure 1A**, and **Methods**). We used immunohistochemistry as a first step to confirm the targeting of Cre to neurons expressing PV, focusing on a set of brain regions enriched with different types of PV neurons (cortical, subcortical, and cerebellum; **Figures 1B, C, S1A, B**). 95.7 ± 0.8% (n = 11858/12368, PV^+^Cre^+^/Cre+) of the Cre-expressing neurons expressed PV across the seven brain regions, confirming high specificity. Immunohistochemistry also confirmed highly efficient targeting of Cre to the PV populations, with 93.6 ± 1.4 % (n = 11858/12641, PV^+^Cre^+^/PV^+^) of the analyzed PV neurons expressing Cre; **Figure 1C**). The Cre expression was specific and efficient throughout the cortical layers (**Figure S1C**). PV is a calcium-binding protein, and its expression varies across neurons and brain regions in an activity-dependent manner^24,25^. This biological variable together with the need for antigen retrieval for Cre detection likely contributed to an underestimation of the overlap between PV and Cre expression. To address functional Cre recombination an adeno-associated virus (AAV) with Cre-dependent expression of ChR2-mCherry was injected into the mPFC of adult PV-Cre rats (N = 3). This confirmed efficient recombination in PV neurons (94.34 ± 0.79% (n = 742/790, PV^+^/mCherry^+^) of the opsin-expressing neurons expressed PV; data not shown but see further below).

**Figure 1.**
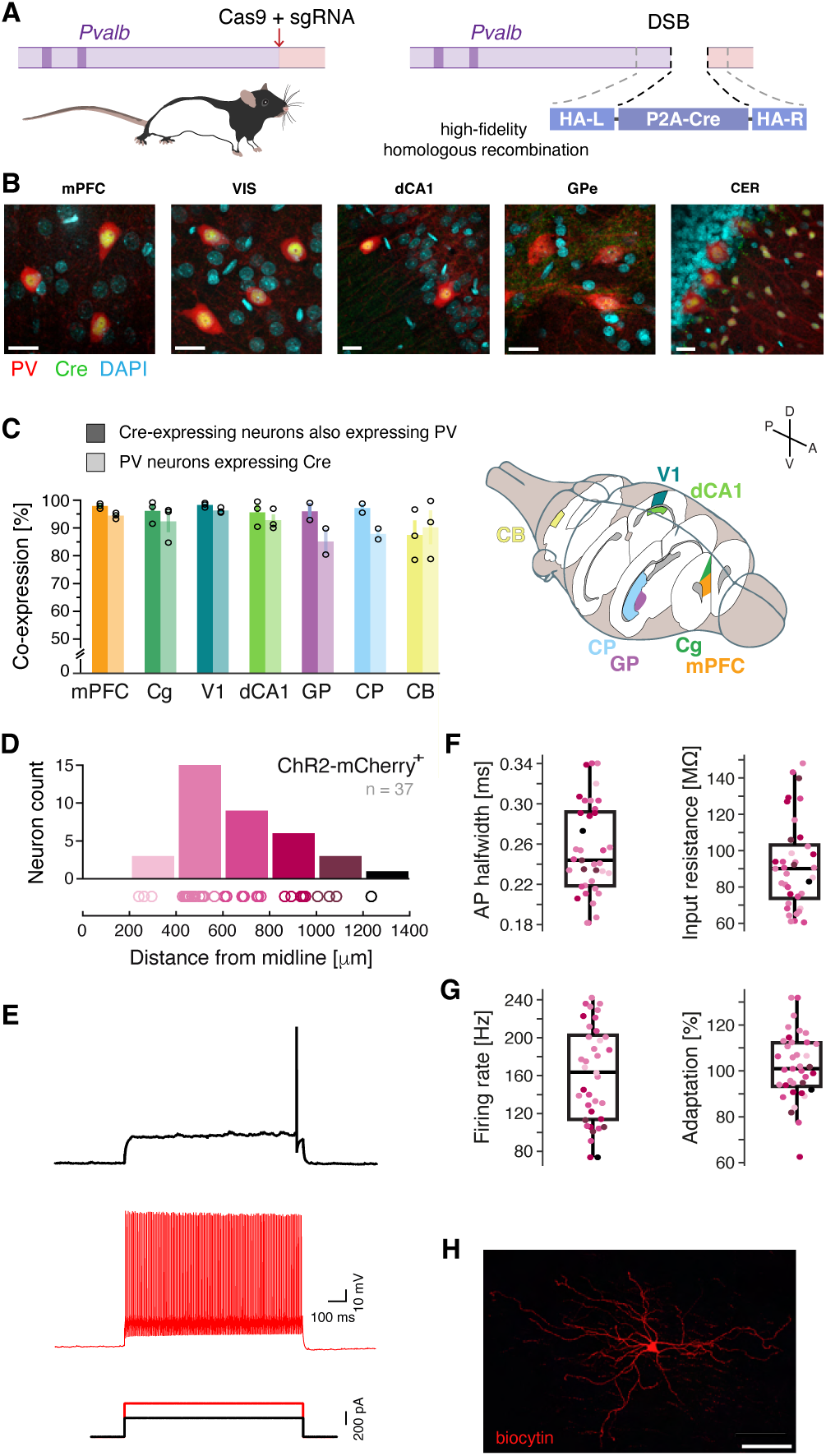
Generation and characterization of the PV-Cre rat line. (**A**) Schematic illustration of the genetic strategy for generation of the PV-Cre knock-in rats. Cre was introduced into the 3’ UTR of the Parvalbumin (Pvalb) locus using a 2A self-cleaving peptide sequence (P2A) between Pvalb and Cre. (**B**) Example of immunohistochemical detection of Cre (green) in PV expressing neurons (red) in the cerebrum and cerebellum of adult PV-Cre rats. Nuclei counterstained with DAPI (cyan). For split channels, see Figure S1B. (**C**) Left: the specificity and efficiency, respectively, of targeting of Cre to PV neurons in seven different brain regions. Dark bars: specificity, i.e., the fraction Cre-expressing neurons also expressing PV: mPFC (PrL + IL): 97.88 ± 0.54 %, Cg: 96.09 ± 2.34%, V1: 98.25 ± 0.49%, dCA1: 95.61 ± 2.69%, GP: 95.92 ± 3.06%, CP: 97.12 ± 1.63%, CB: 87.44 ± 5.23%. Light bars: efficiency, i.e., the fraction PV neurons expressing Cre: mPFC: 94.39 ± 0.63 %, Cg: 92.33 ± 3.81%, V1: 96.25 ± 0.59%, dCA1: 92.75 ± 2.12%, GPe 85.11 ± 4.70%, CP: 87.84 ± 1.94%, CB: 90.20 ± 6.11%. Right: 3D illustration of the brain regions investigated. (**D**) The anatomical location of the recorded ChR2-mCherry expressing neurons, across the cortical depth of the mPFC. Bin size: 200 mm. (**E-G**) Recombined mPFC neurons in adult PV-Cre rats display a typical FS phenotype. (**E**) Response of a recombined mPFC PV neuron to a 1 s current pulse at rheobase current (black) and at twice the rheobase current (red). (**F**) Box plots of the AP half width, and input resistance, respectively, of recombined mPFC PV neurons. (**G**) Box plots of the AP firing rate, and adaptation, respectively, of recombined mPFC PV neurons. (**H**) Representative example of the morphology of a recombined mPFC PV neuron filled with Biocytin. Scale bars: 20 μm (B), 100 μm (G); mean ± SEM (C); boxplots (E, F): black bar: median, box: 25th-75th percentile, whiskers: 99.3% data coverage. Cas9: CRISPR associated protein 9; sgRNA: single guide RNA; DSB: double strand break; HA-L/HA-R: left and right homology arm, respectively; P2A: self-cleaving peptide; mPFC: medial prefrontal cortex; CB: cerebellum; Cg: cingulate cortex area; CP: caudate putamen (striatum); dCA1: dorsal CA1; GP: globus pallidus; V1: primary visual cortex. See also **Figure S1.**

PV neurons in the PFC of adult rats, mice, and monkeys display a typical FS phenotype with narrow APs and minor or no adaptation^26,27^. To characterize the intrinsic electrophysiological properties of the genetically labeled PV neurons in PV-Cre rats, an AAV with Cre-dependent expression of ChR2-mCherry was injected into the mPFC (N = 8 PV-Cre rats), and whole-cell patch-clamp electrophysiology of the labelled neurons (n = 37) was conducted *ex vivo* (**Figure 1D**). In line with previous reports, rheobase current injections elicited narrow APs (0.25 ± 0.05 ms), with a large and short afterhyperpolarization typical of inhibitory FS interneurons (**Figures 1E, F**). The input resistance (92 ± 24 MΩ; **Figure 1F**) was low, similar to the reported values for cortical FS neurons in mice^27^. An AP train evoked at twice the rheobase current revealed typical FS characteristics, with a high AP firing rate (FR; 161 ± 51 Hz) and limited adaptation (102 ± 15%) in response to a 1 second (s) current pulse (**Figures 1E, G**), which is in accordance with earlier studies in both mice and rats^26,27^. All other intrinsic electrophysiological properties confirmed the FS nature of the recorded neurons, including a short membrane time constant and a fast AP downstroke (**Figure S1D**). In addition, neurons labeled with biocytin (n = 6) displayed morphologies typical of cortical PV-expressing basket cells, with multipolar neurites^28^ (**Figure 1H**). No significant laminar differences were observed across the recorded population (**Figures 1F, G, S1D**). Based on the above, we in the current study refer to the inhibitory mPFC interneurons expressing PV as FS-PV neurons.

### Optotagging of FS-PV neurons

To characterize mPFC neuronal and network activities associated spatial processing we paired chronic extracellular recordings with optotagging of FS-PV neurons^29,30^ in a cohort of adult PV-Cre rats (N = 10) engaging in a self-paced reward-seeking task^29,31^. To render mPFC FS-PV neurons light-sensitive, an AAV with Cre-dependent expression of ChR2-mCherry was injected into the mPFC (**Figures 2A, B**). Behavioral training (**Methods**) started one week after viral injection. After the rats had properly learned the task (>50 trials in 1h for three consecutive days) a microdrive holding four movable tetrodes and an optical fiber was implanted (**Figures S2A, B**). While all rats could perform the task after the implantation, detailed post-hoc analysis of the behavior warranted exclusion of three rats, yielding a final data set with extracellular recordings from 97 behavioral sessions (N = 7 PV-Cre rats; **Figures S2C, D** and **Methods**). Optotagging of FS-PV neurons was conducted directly before a subset of the sessions (n = 35). The optotagging and all other optogenetic experiments employed 473 nm blue light (5-10 mW), 1-3 ms light pulses for 1 s, 5-10 light applications. A total of 349 units were recorded and we used a gaussian mixture model (GMM) to cluster the recorded units into WS or NS units, based on the spike width and peak-to-valley ratio^32^ (**Figures S2E, F**). This yielded 281 WS, 54 NS, and 14 unclassified units, respectively, corresponding to ∼80% WS, putative pyramidal neurons, and ∼15% NS, putative inhibitory interneurons (**Figure 2C**) which is in agreement with earlier studies of the rat PFC ^33^. 13 NS units responded with AP firing in direct response to light application (**Figures S2G**), identifying them as FS-PV neurons. Using stimulus-associated spike latency test (SALT) in combination with a spike-shape correlation measure^29^, we confirmed that the 13 optotagged units were directly light-driven (**Figures S2H, I**). The FS-PV neurons displayed a narrow spike width, low peak-to-valley ratio, and high FR, paralleling defining features of cortical inhibitory FS-PV interneurons in mice^30^, and as in mice, light activation of the FS-PV neurons resulted in inhibition of concurrently recorded WS units in the local network (**Figure S2J**). Some NS neurons displayed waveforms similar to the optotagged FS-PV neurons, presumably reflecting non-tagged FS-PV neurons (**Figure S2K**). A substantial fraction of the NS population should be constituted by somatostatin-expressing interneurons^34^.

**Figure 2.**
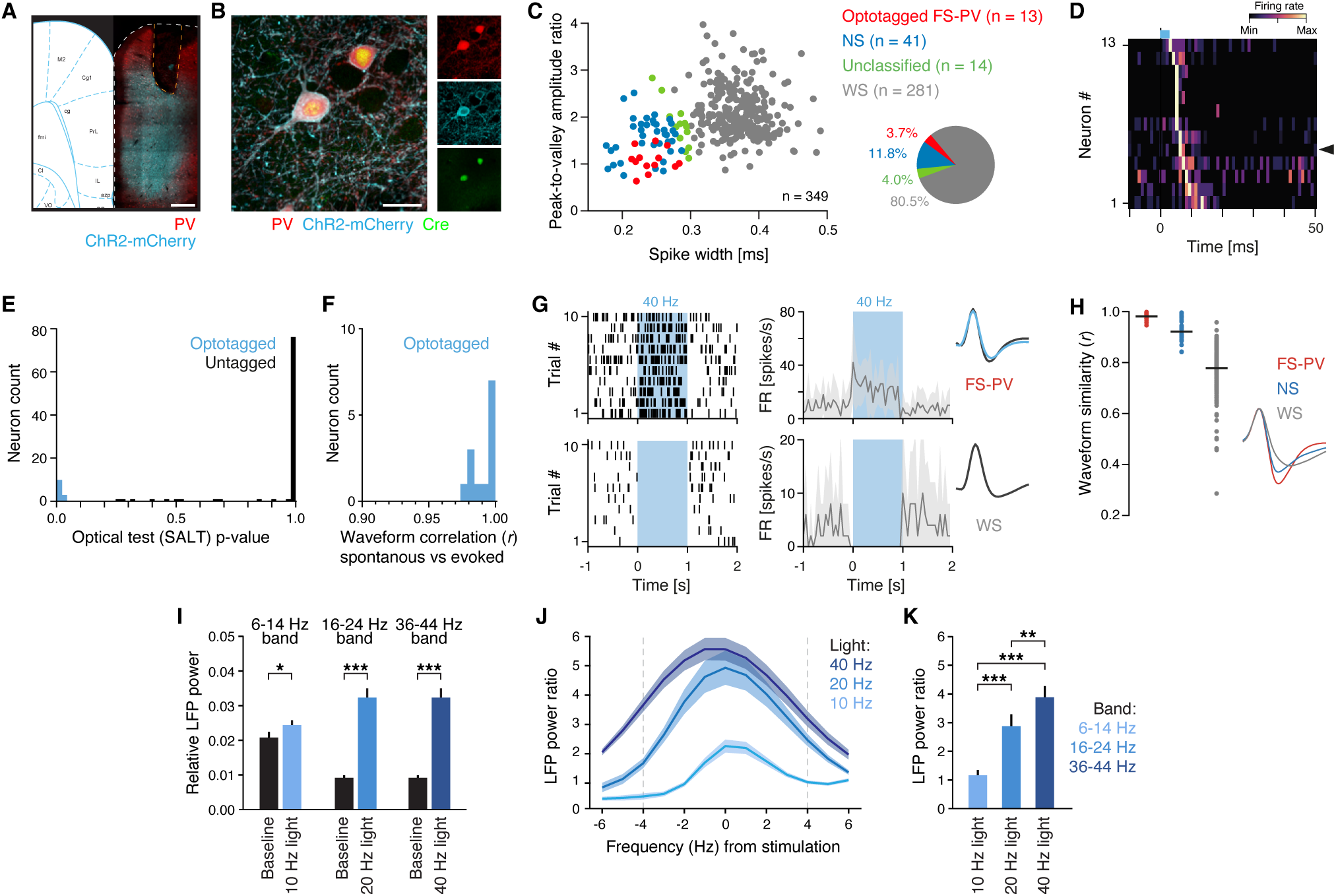
mPFC FS-PV neurons in adult rats generate gamma oscillations. (**A**) Coronal section from PV-Cre rat with ChR2-mCherry expression (cyan) in mPFC PV neurons (red). Orange dashed line: outline of the position of optical fiber. (**B**) Close-up of recombined mPFC PV neurons (red) with ChR2-mCherry expression (cyan) and nuclear expression of Cre (green). (**C**) Left: Plotting and classification of recorded mPFC neurons (n = 349) based on spike waveform properties (spike width, and peak-to-valley ratio (PVR), respectively). Three neuronal cell types were classified: WS (grey, n = 281): spike width: 372 ± 35 μs, PVR: 2.1 ± 0.5, FR: 3.6 ± 3.2 Hz, NS (blue, n = 41): spike width: 239 ± 30 μs, PVR: 1.6 ± 0.4, FR: 4.3 ± 6.0 Hz, FS-PV (red, n = 13): spike width: 242 ± 23 μs, PVR: 1.1 ± 0.3, FR: 13.0 ± 6.6 Hz. The light-activated neurons (red, n = 13) display AP characteristics typical of inhibitory FS-PV interneurons, with short spike width and a low PVR. Right: pie chart with the proportions of classified neurons. (**D**) Averaged and normalized (Min-Max) PSTH aligned to light (3 ms; blue bar) onset for all FS-PV neurons (n = 13). Black arrowhead: the FS-PV neuron in (G). (**E**) Histogram of SALT p-values, identifying direct light-activation (optotagging) of 13 neurons (blue, p < 0.05; n = 102 analyzed neurons). (**F**) There is a high correlation (*r* > 0.95) between the waveforms of light-evoked and spontaneous spikes of the optotagged neurons (n = 13; blue). (**G**) Top row: spike raster of an example optotagged FS-PV neuron responding with increased spiking to 40 Hz blue light (473 nm, 1 s, 3 ms light pulses, left), and the mean FR over 10 trials (right). Bottom row: spike raster of a concurrently recorded WS neuron showing decreased spiking in response to light-activation of FS-PV neurons (left), and the mean modulation over 10 trials (right). (**H**) Left: comparison of the light-evoked waveform (mean) to the spontaneous FS-PV (red), NS (blue), and WS (gray) waveforms, respectively. One dot = one neuron; black horizontal line: mean. Right: average spontaneous waveform of the three neuron types. (**I**) The LFP resonance in response to activation of mPFC FS-PV neurons at 10, 20 or 40 Hz, respectively. Comparison of the relative LFP power at baseline (black) and in response to light application (blues) in the band ± 4 Hz the stimulation frequency. (**J**) Relative LFP power ratio (mean; *P*_light_/*P*_baseline_) in response to activation of mPFC FS-PV neurons at 10, 20 or 40 Hz, respectively. Gray dashed lines: ± 4 Hz the stimulation frequency, i.e., the frequency bands analyzed in (I) and (K). (**K**) Relative LFP power ratio (mean) in the band ± 4 Hz the stimulation frequency in response to activation of mPFC FS-PV neurons at 10, 20 or 40 Hz, respectively. Activation of the mPFC FS-PV neurons at 40 Hz results in significantly more LFP resonance than activation at 10 or 20 Hz. Scale bars: 500 μm (A), 20 μm (B); mean ± SEM (G, I-K); *p < 0.05, **p <0.01, ***p < 0.001. In (I, K) we adjusted the p-values using the Bonferroni correction procedure. See also **Figure S2**.

### FS-PV activation in the rat mPFC generates gamma oscillations

We previously showed that activation of cortical FS-PV neurons in the mouse barrel cortex selectively amplifies oscillations in the gamma band^35^. To investigate how activation of FS-PV neurons in the mPFC of rats resonate in the LFP, we in a subset of the optotagging sessions (n = 22, N = 3 PV-Cre rats) activated the FS-PV neurons with 10, 20, and 40 Hz blue light, respectively. All three stimulations induced LFP resonance around (± 4 Hz) the respective stimulation frequency (**Figures 2I, J)** and also harmonics occurring in integer multiplications of the stimulated frequency (**Figures S2F-H**). To compare the LFP resonance evoked by the different stimulation frequencies we computed the LFP power ratio (Power_light_/Power_baseline_ at the stimulation frequency ± 4 Hz) and found that 40 Hz stimulation of FS-PV neurons resulted in significantly more LFP resonance than the activation at lower frequencies, consistent with specific resonance in the gamma range at the network level^35–37^ (**Figure 2K**).

### Encoding of spatial, movement, and task variables of in the rat PrL

The behavioral setup used for the self-paced reward-seeking task consisted of two platforms connected by a linear track; one platform held a reward spout (reward platform) and for the reward to be delivered the rat needed to visit the other platform (the trigger platform; **Figure 3A**). The task design provides the opportunity to study single trials holding mirrored track traversals where the context and goal of the traversals differ – traversals ‘to reward’ have reward consumption as the goal, whereas traversals ‘to trigger’ have the goal to trigger reward availability. In essence, the reward status differs in the traversals - the reward is a goal in the traversals to the reward platform, and a past event in traversals to the trigger platform.

**Figure 3.**
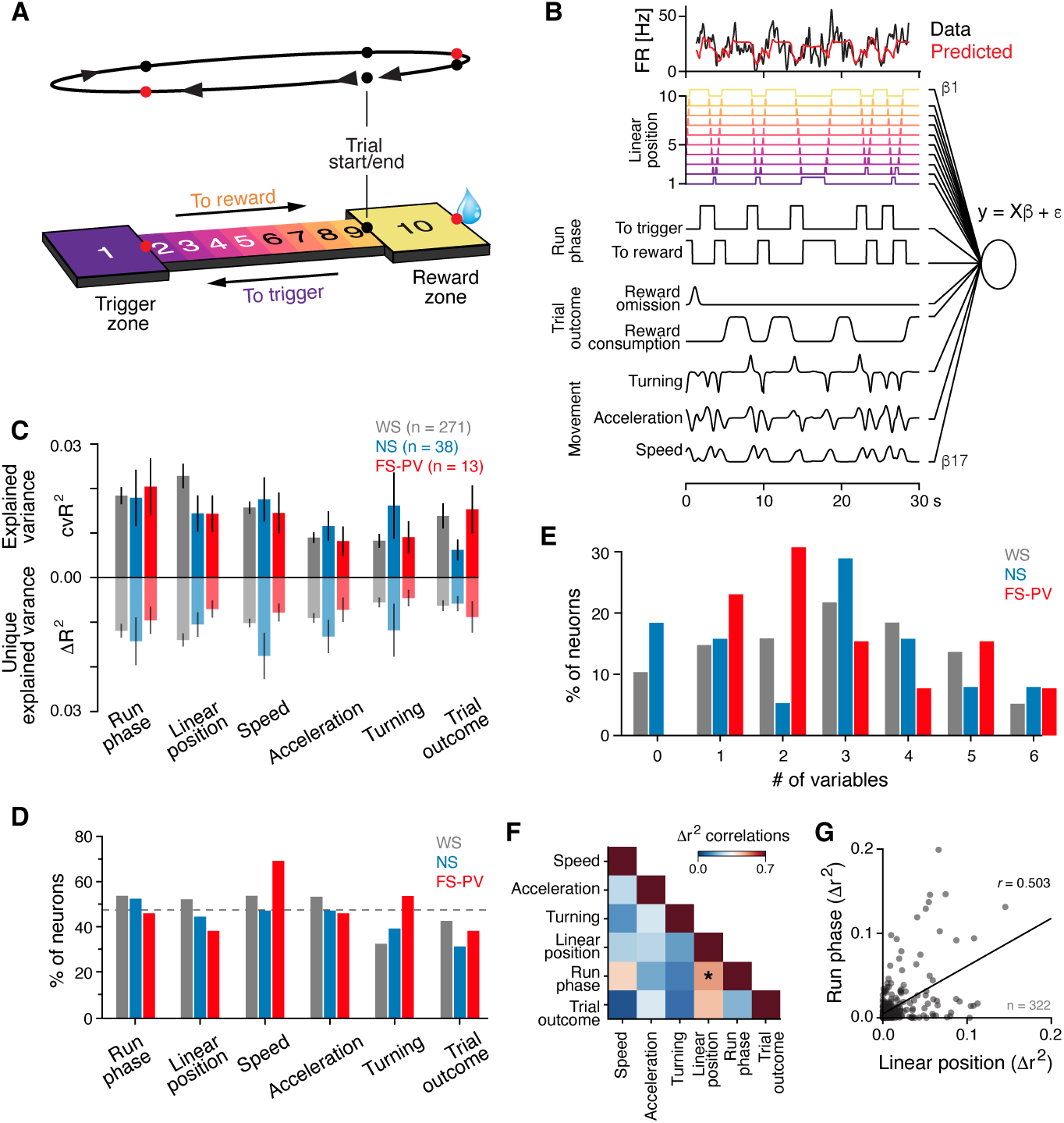
GLMs reveal conjunct encoding of behavioral and task variables in single PrL neurons. (**A**) Schematic of the self-paced reward-seeking task. Top: outline of a single trial, with indication of the traversal direction (arrow heads), and timestamps (colored dots). Red dots: beam break detectors; black dots: timestamps extracted from video. Bottom: illustration of the track with definition of the 10 spatial locations (bins) and indication of the traversal directions. In each trial the rat must visit the trigger zone in order for reward to be delivered in the reward zone. (**B**) Schematic of the GLM approach used to quantify the individual neurons’ encoding of behavioral and task variables. Top: activity of an example neuron (black) and the activity predicted by the full model (red). Bottom: scoring of the behavioral and task variables included in the full model. (**C-E**) Quantification of the correlation between neuronal activity and behavioral and task variables. (**C**) Encoding of model variables by individual neurons were quantified using two approaches. Top: average explained variance (cvR^2^, 5-fold cross-validated) of single variable models. Bottom: average unique explained variance (Δr^2^) of the variables. (**D**) The fractions neurons with significant encoding (p < 0.01, permutation test) of the different behavioral and task variables. Dashed line: mean across all variables (47 ± 4.4 %). (**E**) Number of variables that are significantly encoded by each neuron. This revealed that many neurons encode more than one variable. (**F**) Pearson’s correlation between the unique explained variance (Δr^2^) for all combinations of variables. There is a significant correlation between the animal’s position on the track (*linear position*) and the direction of the track traversal (*run phase*; p < 0.05, Holm-Bonferroni correction as compared to shuffled data). (**G**) Scatter plot of the unique explained variance (Δr^2^) for the *linear position* (x-axis) vs the *run phase* (y-axis) for all individual neurons (n = 322), revealing a significant correlation. Mean ± SEM (C). See also **Figure S3**.

To quantify neuronal encoding, we used generalized linear models (GLMs) to fit the individual neuron’s activity with behavioral and task variables as regressors (**Figure 3B**)^38–40^. Both categorical variables (linear position (in one of 10 spatial bins in the behavioral setup), run phase (to reward or to trigger (**Figure S3A**)), trial outcome, trial time, trial type, trial history), and continuous variables (speed, acceleration, turning) were included (**Supplementary Movie**, and **Methods**). The performance of the models, reflecting neuronal encoding, was quantified by calculation of the explained variance (cvR^2^; squared Pearson’s correlation (r^2^) between the predicted and actual test data), obtained by 5-fold cross-validation. As a first step we identified which variables were encoded by the PrL neuronal population and therefore should be included in the models. To this end, we fitted the activity of the individual neurons using models that incorporated only a single variable and we thereafter ranked the variables based on their ability to predict the neuronal activity (**Figure S3B**). The variables were subsequently added one by one, in descending order, until all variables were included in the models. This revealed a lack of significant encoding of trial time, trial history, and trial type, as the inclusion of these variables did not significantly improve model performance – these variables were therefore not included in the final (full) models (**Figure S3C**). Across all recorded neurons the animals’ linear position was the variable best predicting neuronal activity (2.5 ± 0.50 %), followed by the run phase (2.0 ± 0.37 %), and the running speed (1.6 ± 0.26 %; **Figure S3B**). On average, the full models encompassing six variables explained 7.0 ± 1.1% (95% CI) of the variance (**Figure S3C**), comparable to previous reports^40,41^.

We observed that the explained variance of the full models was lower than predicted (**Figure S3C**), indicating an overlap in the predictive power of some variables, i.e., certain variables contained redundant information. This suggests that cvR^2^ is a suboptimal measure of variable encoding. To optimally capture each variable’s unique contribution (Δ*R*^2^) to the models we created reduced models, by removing single variables and quantified the reduction in cvR^2^ relative to the full models (**Figure S3D)**^38–40^. As expected, the removal of any of the variables reduced the explained variance (cvR^2^) of the full models, with linear position, the run phase, and the running speed, causing the largest reductions (**Figure S3D, E**). Overall, the animal’s linear position was the best predictor of the activity of the PrL neurons (**Figures S3B, E**).

### Cell type-specific encoding of behavioral and task variables in the rat PrL

Our observations of spatial and movement encoding in the PrL corroborate previous findings in mice^5,40^ and rats^42^. However, to the best of our knowledge, the activities of FS-PV neurons have not been accounted for. We therefore as the next step separately characterized the encoding by the FS-PV, WS, and NS populations. Both the explained variance (cvR^2^) of the single variable models, and the unique contribution (ΔR^2^) derived from the reduced models indicated a lack of significant differences in the encoding strength of the individual variables between the three neuronal populations (p > 0.05; t-test, **Figure 3C**). Comparable fractions of WS, NS, and FS-PV neurons encoded the individual variables, and overall, the behavioral and task variables were significantly encoded by 47 ± 4.4 % of the PrL neurons (**Figure 3D**). A few WS, and NS, neurons did not encode any of the variables (WS: 28/271; 10.3%, NS: 7/38; 18.4%; **Figure 3E**). The majority of the neurons (238/322; 73.9%) encoded more than one variable, indicative of mixed selectivity^43,44^. To investigate this further we computed the pair-wise correlation between the encoding strength of all variables, which revealed a significant linear relationship between the unique contribution (ΔR^2^) of the animal’s linear position on the track and the run phase (p < 0.05 permutation test, Holm-Bonferroni correction; **Figures 3F, G**). This indicates that subpopulations of PrL neurons encode the rat’s spatial location on the track and also the direction of the track traversal, suggesting that neurons in the PrL display context-dependent spatial representations (see further below).

### Context-dependent spatial encoding in the rat PrL

The GLM revealed that spatial representations in the rat PrL were modulated by the context of the track traversal. To characterize how the contexts of the two traversals were represented in the neuronal activity, we analyzed the firing patterns in relation to both the animal’s spatial location and the direction of the track traversal. This identified sequential firing along the track’s 10 bins, with overall one population firing in one of the two directions, and a different population in the other direction (**Figure S4A**). Comparison of the activities between odd and even trials revealed that this pattern was consistent (spatial consistency) and did not reflect random fluctuations in the activity (**Figure S4A-C**)^5,40^. The tiled firing was particularly evident in the large WS population, with individual WS neurons firing in only a small range of spatial bins in one of the directions (**Figures 4A, B, S4C**). Hence, the WS neurons carried information not only about the animal’s spatial position on the track but also about the context of the track traversal. The spatial context was also evident in the activities of the FS-PV population, reflected by significantly higher activity in traversals to the trigger zone than in traversals to the reward zone (p < 0.05, t-test; **Figures 4A, C, D, S4C**). The activity was particularly differential when the animal reached the respective platform, with >60% of the FS-PV neurons displaying increased activity in the trigger zone, and no FS-PV neurons showing increased activity in the reward zone (**Figures 4C, E**). Quantification of the similarity of the individual neurons’ FR in the two traversals as expected revealed weak correlations (Pearson’s (*r)*), especially in the later part of the traversals (i.e., closer to the platforms), and particularly for the FS-PV neurons (**Figure 4F**). Across the three cell types 63.0 % (n = 201/319) of the neurons displayed significantly different mean activities in traversals to reward vs traversals to trigger (WS: 64.2%, n = 172/268, NS: 55.3%, n = 21/38, FS-PV: 61.5%, n = 8/13; **Figure 4D**). To quantify the spatially related activity in the respective traversal, we computed the spatial information (SI)^5^ from spatially binned firing rates (**Methods**). This identified higher SI in traversals to reward than in traversals to trigger for all three cell types (**Figure 4G)**, i.e., the animals’ position in space (linear position on the track) was more prominently represented during traversals to reward than during traversals to the trigger.

**Figure 4.**
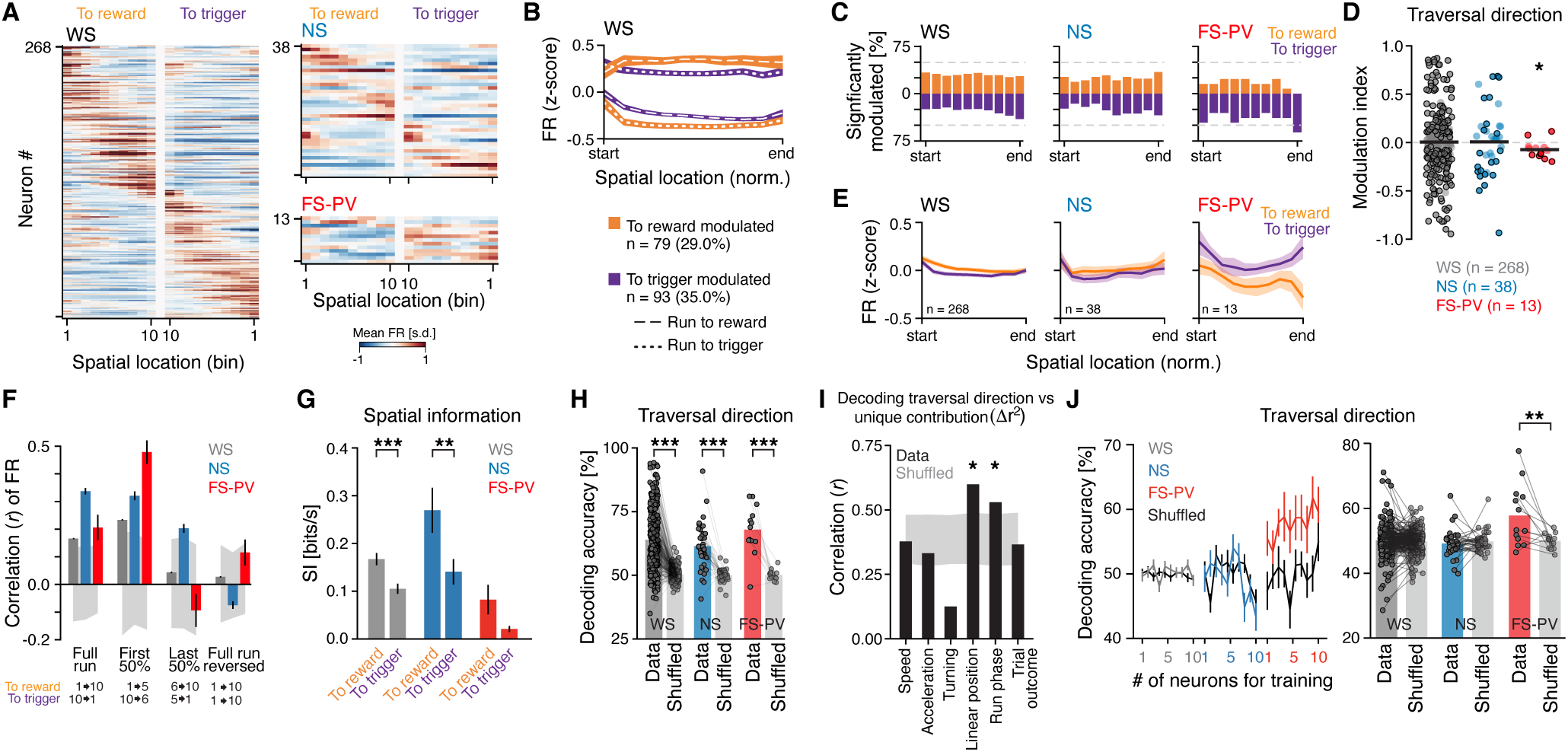
Encoding of spatial context. (**A**) The trial-averaged FR (z-scored) of all individual neurons, split by cell type and traversal direction. The neurons are peak sorted by the FR in traversals to reward and plotted in the same order for both traversal directions. (**B**) The mean activity (z-scored) of WS neurons significantly modulated by the animal’s traversal direction (p < 0.05, Wilcoxon rank-sum test). Different WS populations are activated in traversals to reward vs to trigger. The two populations show opposing activity patterns during the respective traversals. (**C**) Fraction of neurons with significantly modulated activity (p < 0.05, permutation test) at each of the 10 track positions (bins) in the two track traversals. Orange: increased activity in traversals to reward; purple: increased activity in traversals to trigger. Gray dashed lines: 50%. (**D**) The modulation index (MI) of the traversal direction for the three cell types. Positive values: increased activity in traversals to reward, negative values: increased activity in traversals to trigger. Mean MI (black line): WS, n = 268: 0.008 ± 0.32, p = 0.68; NS, n = 38: 0.027 ± 0.309, p = 0.61; FS-PV, n = 13: -0.068 ± 0.086, p = 0.018; t-test. The FS-PV population has significantly higher activity in traversals to trigger than in traversals to reward. Black circles; significantly modulated neurons (p < 0.05, Wilcoxon rank-sum test; to trigger: WS: n = 93, NS: n = 13, FS-PV: n = 6; to reward: WS: n = 79, NS: n = 8, FS-PV: n = 2). (**E**) The mean activity (z-scored) of the three cell types in the two traversal directions. The FS-PV population displays higher activity in traversals to the trigger than in traversal to the reward, with particularly dissimilar activities in the end of the traversals (i.e., when the animals reach the respective platform). (**F**) Comparison of the FR in the two traversal directions. The correlation (*r*) between the FR in traversals to reward vs to trigger was computed for each neuron (numbers as in (D)). The FR of the respective cell type was most similar in the first half of the traversals, and least similar at the end of the traversals, and this was particularly true for the FS-PV neurons. First 50%: WS: 0.23 ± 0.002, NS: 0.34 ± 0.017, FS-PV: 0.48 ± 0.043; Full run: WS: 0.17 ± 0.002, NS: 0.29 ± 0.013, FS-PV: 0.21 ± 0.046), Last 50%: WS: 0.05 ± 0.003, NS: 0.15 ± 0.017, PV: -0.09 ± 0.059; Full run reversed: WS: 0.0 ± 0.002, NS: -0.1 ± 0.015 and PV: 0.12 ± 0.047. Grey area: 95% confidence interval of shuffled data. (**G**) The spatial information (SI) was consistently higher in traversals to reward than in traversals to trigger, with significant differences for the WS neurons and the NS neurons, respectively. WS: p = 1.7×10^-6,^ NS: p = 0.003, FS-PV: p = 0.054, paired t-test. Colors and numbers as in (D). (**H**) For all three cell types, the traversal direction could be predicted based on the neuronal activity, above chance level (WS: p = 6.0×10^-58^, NS: p = 7.5×10^-9^, PV: p = 2.8×10^-5^, paired t-test). This indicates that all three neuronal populations encode the traversal direction. One dot = one neuron, numbers as in (D). (**I**) Correlation between the decoding accuracy (**H**) and the unique contribution of the variables used in the full model (Figure 3C), revealing a significant correlation specifically for the *linear position* and the *run phase.* Gray area: 95% confidence interval of shuffled data. (**J**) Left: SVMs were used to predict the animals’ traversal direction from the activity of individual neurons. For this, the SVMs were trained on data from 1-10 randomly selected, different, neurons. Black traces: predictions from SVMs trained with shuffled traversal direction labels. Right: the mean decoding accuracy for all individual neurons, based on the analysis in left. The traversal direction could be predicted above chance level only from the activity of the FS-PV neurons (FS-PV: p = 0.004 WS: p = 0.172, NS: p = 0.705, vs shuffled control data; paired t-test), demonstrating more homogeneous encoding of traversal direction within the FS-PV population than within the WS, or NS, populations. Mean ± SEM (B, E, F, J). *p < 0.05, **p < 0.01, ***p < 0.001. See also **Figure S4**.

Our analyses of context-dependent spatial representations employed trial-averaged activities, which do not account for variations between individual trials, and we therefore next addressed how reliably the spiking of individual neurons on a trial-to-trial basis could predict the direction of individual traversals. For this, we trained linear support vector machines (SVMs) to decode the direction of the traversals, revealing high and significant decoding accuracy for all three cell types compared to shuffled data (WS: 66 ± 1.5 %, NS: 64 ± 3.7%, PV: 69 ± 6.0%; **Figure 4H**). Neurons with strong encoding of the run phase (GLM, **Figure 3C)** would be predicted to display high decoding accuracy for the traversal direction. To confirm this, we analyzed the correlation between the decoding accuracy for the traversal direction and the unique contribution of all variables in the full model. As expected, a significant correlation was found for the run phase, and also the linear position, but not for the other variables in the full model (**Figure 4I**). The significant correlation between the encoding of linear position and the decoding accuracy for the traversal direction provides additional support for conjunct encoding of the animals’ position and the reward location in single PrL neurons.

To inquire how homogeneously spatial context was encoded across the neurons in each cell type we used SVMs to predict the direction of the traversal from the activity of individual neurons. The SVMs were trained on data from 1-10 randomly selected, different, neurons of the same cell type. This revealed that the traversal direction could be predicted above the chance level from the activity of the FS-PV neurons but not from the WS, or the NS, neurons (**Figure 4J**). This indicates that spatial context is reflected by similar activity patterns across the FS-PV population, in line with FS-PV activities being low in one traversal (to reward) and high in the other (to trigger; **Figures 4C-E**), while the WS and NS neurons displayed more tiling activities regardless of direction of the traversal (**Figure 4A**). On the whole our analyses identify neurons in the PrL that do not only encode the animals’ actual location in space (‘place cells’), or a specific location relative to the start or stop of the movement trajectory (i.e., a specific ‘trajectory phase’)^31^ but rather the context of the movement in space, i.e., the spatial context.

### Context encoding in relation to movement variables

Signals related to movement are considered ubiquitous and can confound the correlation of neuronal activity to cognitive and other task-related processes^45,46^. As we observed that PrL neurons encode movement variables (**Figure 3**) we therefore next moved our attention to the animals’ movement on the track (speed, acceleration, and turning) and the encoding of these continuous variables. We found the whole continuum of movement speeds (0 to 100 cm/s) to be reflected in the activity of the WS population, with some WS neurons being tuned to a narrower range of speeds than others (**Figure 5A**). In contrast, the WS activities associated with acceleration were strikingly dichotomized, with absolutely most WS neurons being tuned to either acceleration or deceleration (**Figure 5A**). Also the WS responses to ipsi- vs. contralateral turning on the platforms were dichotomized (**Figure 5A**). Importantly, the animals’ traversals to reward were faster and had a different acceleration pattern than traversals to trigger (**Figure 5B**) and it was therefore a possibility that the neuronal activities interpreted as encoding of spatial context (**Figure 4**) would be better explained by variables related to the animals’ movement on the track. To address this, we used a GLM to quantify the encoding strength of speed, acceleration, and spatial context, respectively, during the animals’ running on the track (**Figure S3A**; **Methods**). Encoding of spatial context was isolated by generation of a new variable, position with direction, which captures both the animal’s position and direction on the track. Reduced models were used for quantification of the unique explained variance of the three respective variables (**Figure S3D; Methods**). This identified that the position with direction variable explained significantly more of the variance than both the speed and acceleration variables only for the FS-PV population (**Figure 5C**), i.e., spatial context was encoded significantly more strongly than the movement variables by only this population. While position with direction was the strongest predictor also for the WS activity, this variable did not explain significantly more of the variance than the speed variable. In essence, while spatial context is encoded by the WS population, some activity differences in the two track traversals can be attributed to speed encoding. In support of this, similar fractions of WS neurons encoded speed and position with direction, respectively (**Figure 5D**). Overall the analyses show that while spatial context is encoded by all three PrL populations, this encoding is particularly pronounced in the FS-PV population.

**Figure 5.**
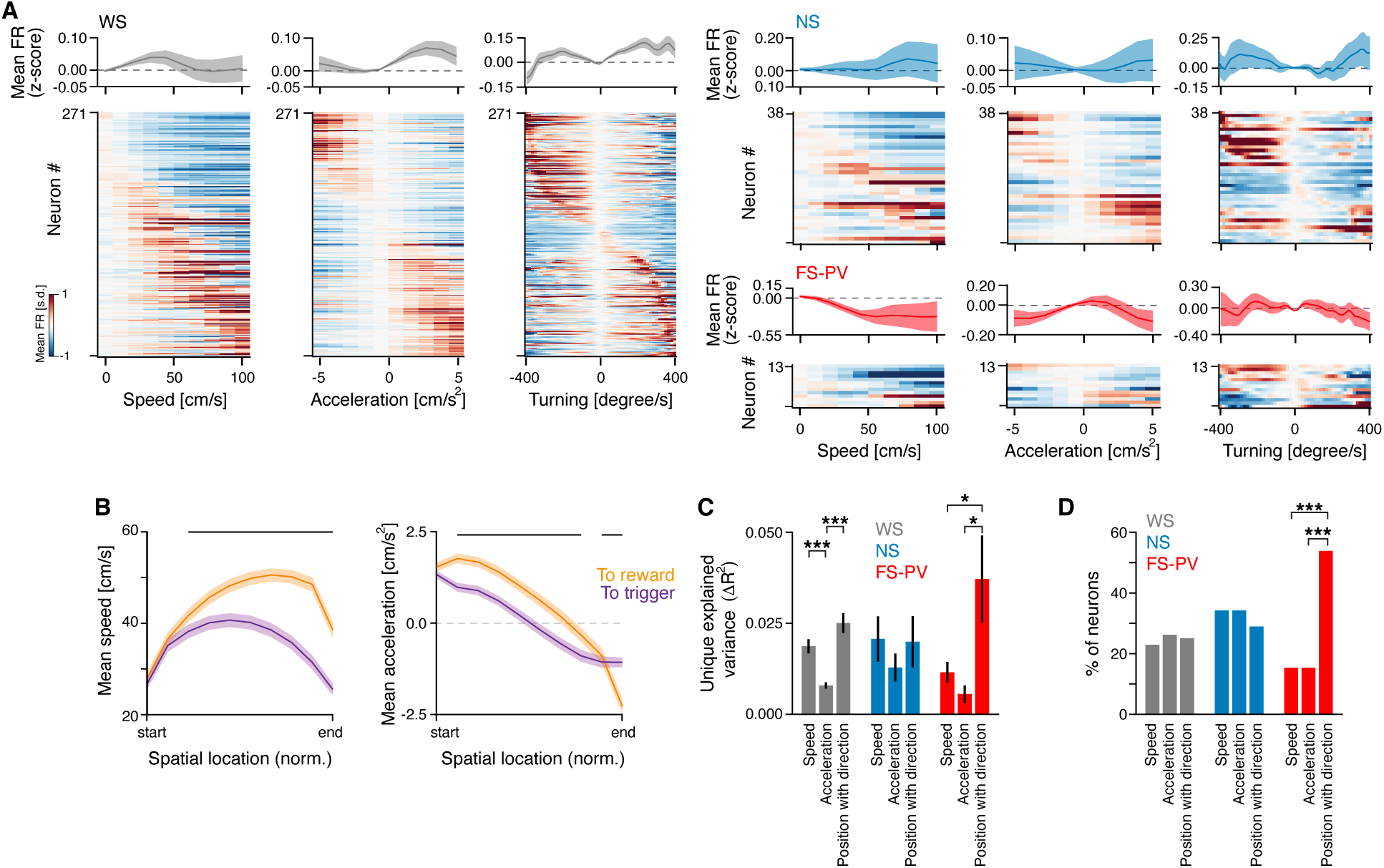
Encoding of spatial context in relation to movement variables. (**A**) Top: mean firing rate (z-scored) as a function of speed (cm/s, bin size = 10 cm/s; left), acceleration (cm/s^2^, bin size = 1 cm/s^2^; middle), and turning (degrees/s - head angular velocity ipsilaterally (positive values) or contralaterally (negative values) to the recorded hemisphere, bin size = 10 degrees/s; right). Bottom: the trial-averaged FR (z-scored) for all individual neurons, split by cell type. The neurons are peak sorted in the individual panels. (**B**) Mean speed (left) and mean acceleration (right) in traversals to reward and trigger, respectively. Black horizontal lines: significant difference between the two traversal directions (p < 0.05, paired t-test). (**C**) The unique explained variance (Δr^2^) of the variables *speed*, *acceleration,* and *position with direction,* respectively. Only the activity of the FS-PV neurons is significantly better explained by *position with direction* than by *speed* or *acceleration*. (**D**) The fraction neurons with significant encoding (p < 0.01, permutation test) of the variables *speed*, *acceleration,* and *position with direction*, respectively. A significantly larger proportion of the FS-PV neurons encode the variable *position with direction* than the variables *speed* and *acceleration,* respectively (p < 0.01, binomial test). Mean ± SEM (A, B, C). *p < 0.05, **p < 0.01, ***p < 0.001; paired t-test (B, C).

### Prefrontal gamma oscillations in relation to spatial processing

LFP oscillations reflect network activity patterns and are ubiquitous signatures of operating circuits. In line with this, oscillations are used for readout of computations and behavioral variables, in the PFC often with a focus on gamma oscillations due to the strong correlation between prefrontal gamma activity and cognition^30,47,48^. Building on these concepts we next investigated the modulation of the LFP activity during the animals’ running on the track. Spectral analysis of the LFP oscillations revealed differences between traversals to reward and traversals to trigger, with the power in the gamma band (30-50 Hz) being significantly higher in traversals to reward than in traversals to trigger (**Figures 6A-C, S5A-C**). Single-trial analysis demonstrated a pattern of bursts of increased gamma power rather than a continuous elevation (**Figures 6D, S5D**), with the rate of gamma bursts being significantly higher in traversals to reward than in traversals to trigger (paired t-test, p < 0.001; **Figure 6E**). In the hippocampus, the expression of gamma oscillations has been correlated to the speed of movement^17,49^ and as the speed differed in the two traversal directions (**Figures 5B, 6F, S5E**) we investigated the relationship between speed and gamma power. We found an overall positive correlation, but importantly, accounting for the differential running speeds in the two directions did not abolish the difference in gamma power between the two traversal directions (**Figures 6G, S5F, G**). Thus, the prefrontal computations involved in our asymmetrical reward-seeking task were reflected by modulation of the gamma band activity.

**Figure 6.**
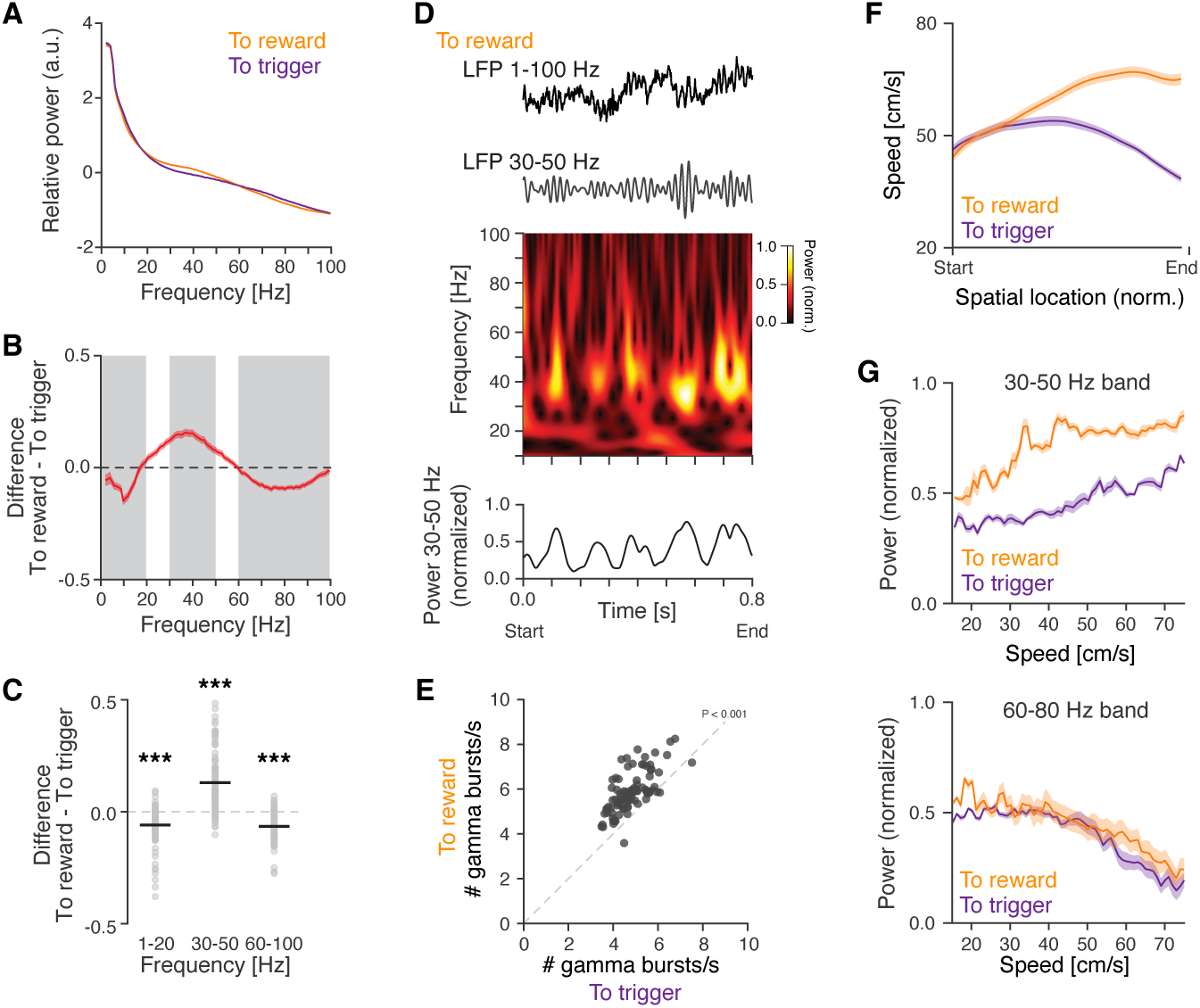
Modulation of PrL gamma oscillations by reward-seeking behavior. (**A-C**) The spatial context is reflected in the LFP, with traversals to reward being characterized by increased gamma oscillations. (**A**) The relative LFP power (1-100 Hz) in traversals to trigger vs to reward. (**B**) The difference in relative power in traversals to trigger vs to reward. Gray bars: frequency bands with statistics in (C). (**C**) The difference in relative power in three frequency bands, between traversals to trigger and traversals to reward. Positive values: increased relative power in traversals to reward, negative values: increased relative power in traversals to trigger. Black lines: mean, gray circles: individual recording sites. (**D-E**) The increased gamma power in traversals to reward is accompanied by an increased rate of gamma bursts. (**D**) Example LFP traces (0.8 s) from a single traversal to reward. From the top: raw LFP; band-pass filtered LFP (30-50 Hz); spectrogram (10-100 Hz); extraction of the mean power in the gamma band (30-50 Hz). Dashed line: 75^th^ percentile power. (**E**) Detection of individual gamma bouts at the 91 recording sites during traversals to reward (y-axis) and to trigger (x-axis), respectively. Burst/s: to reward: 5.9 ± 0.1, to trigger: 4.8 ± 0.1; p < 0.001, paired t-test. (**F-G**) The relationship between running speed and power in the LFP spectrum. (**F**) The running speed and its dynamics differ depending on the direction of the track traversal. (**G**) The LFP power in two frequency bands after adjustment for the differential running behavior in the two track traversals. Top: while the gamma power overall increases with the running speed, the power is higher in traversals to reward than in traversals to trigger. Bottom: the power in the 60-80 Hz band in contrast decreases with the running speed and does not differ depending on the direction of the track traversal. 91 recordings sites (A-E, G), one dot = the mean at one recording site (C, E); Start/end = bin 2 or 9, depending on the traversal direction (see Figure 3A). Mean ± SEM (A, B, F, G). ***p < 0.001.

Spatial encoding in the rat hippocampus is signified by neuronal spiking coupled to the phase of ongoing network oscillations in the gamma and theta bands^50,51^. However, the relationship between local network oscillations and neuronal discharge in prefrontal operations remains to be deciphered, including during spatial processing, and we therefore next investigated the strength of spike phase-coupling during the track traversals. We focused on the gamma band (30-50 Hz) and quantified spike-phase-coupling using two different measures: pairwise phase consistency (PPC^52^) and mean resultant length (MRL), respectively. At the population level, both measures identified increased spike-gamma phase-coupling during traversals to reward for all three PrL cell types, although only the MRL showed statistical differences between the two traversal directions (**Figures 7A, B**). We thereafter investigated a possible relationship between the phase-coupling and the spatial representation by PrL neurons. Due to the low number of FS-PV and NS neurons, respectively, we focused on the WS population. The WS neurons were divided into four groups based on their gamma phase-coupling properties – (A) WS neurons with significant gamma phase-coupling in traversals to reward, (B) in traversals to trigger, (C) in both traversal directions, and WS neurons lacking significant gamma phase-coupling (D; **Figure 7C**). None of the populations displayed significantly different FR in the two traversals (**Figure 7D**). To quantify the spatially related activity in the four subpopulations, we as before (**Figure 4G**) calculated the SI from spatially binned firing rates. This identified that the higher SI in traversals to reward than in traversals to trigger identified for all three cell types (**Figure 4G**) persisted across the four WS populations, i.e., the SI was highest in traversals to reward regardless of the gamma phase-coupling properties of the WS neurons (**Figure 7E)**. This strongly indicates that spike gamma phase-coupling is not an overarching feature of spatial processing in the rat mPFC. In support of this, the levels of SI in traversals to reward, and in traversals to trigger, respectively, were similar across the four WS populations (**Figure 7E**). Furthermore, while the WS population with spike-gamma phase-coupling in both traversal directions showed a positive correlation between the PPC and the SI, this correlation was weak and non-significant (**Figures 7F, G**). Two informative observations can be made from the data. First, the spatial information in the firing of the PrL neurons was consistently increased when the animals moved towards an expected reward (across the cell types; **Figure 4G**), regardless of the relationship between the spiking and the local gamma oscillations (WS neurons; **Figure 7E**). Second, subpopulations of WS neurons displayed gamma phase-coupling selectively in one of the traversal directions, although the neurons’ SI was high in both directions. Both observations identify that the spatial representation in the WS neurons is not related to their gamma phase-coupled firing. This further suggests that while gamma phase-coupling is not directly related to spatial encoding in the mPFC, gamma phase-coupling is a feature of discrete WS ensembles during goal-directed reward seeking behavior. It remains to be established what operations the gamma phase-coupling support, and what computations the modulations of the gamma oscillations reflect (**Figure 6**).

**Figure 7.**
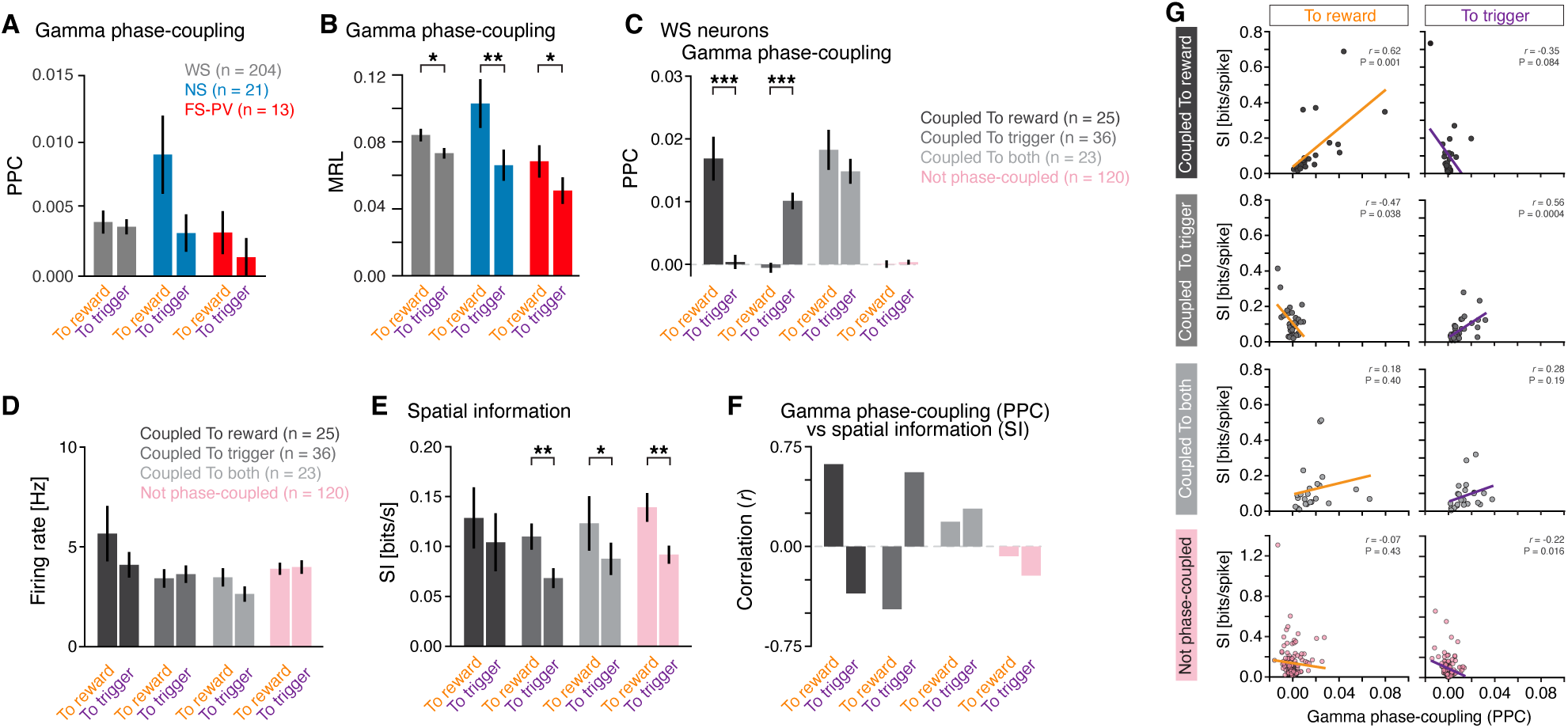
Spike-gamma phase-coupling of PrL WS neurons do not reflect spatial processing. (**A**) No significant differences in the strength of the spike-gamma phase-coupling (quantified as pairwise phase consistency, PPC) were found between the two traversal directions (WS: p = 0.720, NS: p = 0.061, FS-PV: p = 0.119). (**B**) The strength of spike-gamma phase-coupling in the two traversal directions, quantified as MRL. This measure identifies that the spiking of neurons of all three cell types is significantly more phase-coupled to local gamma oscillations (30-50 Hz) in traversals to reward than in traversals to trigger (WS: p = 0.01, NS: p = 0.003, FS-PV: p = 0.04). (**C-G**) Lack of support for gamma phase-coupling of WS neurons being a feature of spatial encoding in the PrL. (**C**) WS neurons were divided into four groups, based on their gamma phase-coupling properties. (**D**) None of the four populations of WS neurons displayed significantly different FR in the two traversals. (**E**) Across the four WS populations, the SI was highest in traversals to reward, regardless of the neurons’ spike-gamma phase-coupling. (Locked to reward: p = 0.42, Locked to trigger: p = 0.04, Locked to both: p = 0.04, not locked: p = 0.001). SI to reward did not differ between the four WS groups (p = 0.75, ANOVA) (**F-G**) Analysis of the relationship (Pearson’s correlation (r)) between the spike-gamma phase-coupling (PPC) and the spatial representation (SI) in the two traversal directions. (**F**) Bar plots of the correlation between PPC and SI in the two traversal directions. (**G**) Scatter plots of the PPC (x-axes) and SI (y-axes), for all four WS groups in both traversal directions. Mean ± SEM (A-E). *p < 0.05, **p < 0.01, ***p < 0.001; paired t-test (A-E).

## Discussion

We here introduce a transgenic rat line with faithful and specific Cre expression in PV neurons and demonstrate its utility for cell-type specific experiments *ex vivo,* and *in vivo* during behavior, respectively. Our *ex vivo* experiments demonstrate that mPFC PV neurons in adult rats display properties typical of inhibitory FS interneurons and in many ways mirror cortical FS-PV neurons in adult mice. The optogenetic experiments demonstrated direct light-activation of mPFC FS-PV neurons and concurrent silencing of WS, putative pyramidal, neurons in the local circuit.

The theoretical and experimental support for the importance of FS-PV activities in the generation of gamma oscillations is ample, and optogenetics has been frequently used in mouse studies to address how gamma rhythmicity relates to information processing and cognition in both health and disease^35–37,48^. Optogenetic drive of FS-PV neurons in the rat mPFC resulted in specific resonance in the gamma range at the network level, replicating findings in the adult mouse cortex^35,36,53^. Optogenetic manipulation of PV neurons in different brain regions has the past 15 years found utilities beyond studies of the neurons *per se* - photostimulation of cortical FS-PV neurons has proven a powerful means for rapid inactivation of cortical microcircuits in mice^54,55^ and gamma frequency activation of FS-PV neurons in cortex or hippocampus has been used for both restoration and improvement of cognition and/or behavior^30,48,56^. Our data suggests that the PV-Cre rats open up for new research approaches also in this species.

### Encoding of spatial context by mPFC FS-PV neurons

mPFC neurons tend to be less spatially selective than hippocampal place cells^5^, and the spatial processing is intricately tied to a variety of factors beyond mere spatial location, including goal locations, past choices, the surrounding environment’s context, and task structures^7–10^. Thus, the mPFC appears to emphasize a location in relation to cognitive and behavioral variables relevant to the task at hand. Our study adds to this body of work and demonstrates that while different cell types in the rat PrL encode discrete variables reflecting the animal’s spatial location, movement (speed, acceleration, turning), and if the behavior was rewarded or not (trial outcome), subpopulations of single neurons encode the animal’s spatial position in relation to a rewarded goal. While we identified the encoding strength of discrete behavioral variables to be similar across the three cell types studied (**Figure 3**), the spatial context was particularly encoded by the FS-PV population (**Figure 5**). A recent study identified that neurons in the mouse anterior cingulate cortex are tuned to a specific trajectory phase, i.e., a spatial location relative to the start and end of a traversal on a linear track, regardless of the animal’s direction on the track^31^. Importantly, in this study, the animals could collect a reward on both ends of the track, while in our study reward was only available in one of the platforms. Together the studies suggest that the asymmetric presence of a reward site is what drives the differential spatial encoding of the two track traversals in our study. Supporting this, recent studies using a spatial working memory task have shown that prefrontal pyramidal firing is more influenced by the animal’s direction (inbound vs. outbound in relation to reward) than by the task phase (sample vs. choice)^7,40^.

We found that the spiking of all three cell types held more SI during traversals to reward than during traversals to trigger, suggesting stronger spatial representation when the rats ran towards the reward. In line with this, the spatial information in the spiking of putative pyramidal neurons in the PFC has been found to be higher when mice approach a reward site compared to when moving away from it, a difference that was abolished when the reward was removed^7^. We show here that the traversals to the reward site are associated with increased power and bursts of gamma oscillations in the PrL, i.e., the asymmetric reward-seeking is associated with modulation of the LFP activity in the gamma band. As gamma rhythms are suggested to temporally organize spiking, we enquired about the relationship between the WS spiking and the gamma cycle phases, and if spike-gamma phase-coupling was a feature of spatial processing in the PrL. Our results do not support that local gamma oscillations organize spiking activities part of spatial encoding in the PrL and instead suggest that the strong modulation of the gamma band activity is related to other prefrontal operations during the reward seeking behavior. Future work could employ a longer linear track for ambulating traversals, optimized for investigation of spike-phase relationships to lower LFP frequencies, including theta frequencies. The generated PV-Cre line presents opportunities for detailed enquiries in rats about the role of inhibitory PV neurons in population activities and their relationship to theta and gamma rhythms, across the cortex and hippocampal formation.

## Author contribution

HB performed all behavioral experiments, analysed electrophysiological, optogenetic, histological, and behavioral data, and plotted the data. HK contributed to surgeries and experimental planning, and analyzed and plotted LFP data. SÄR performed histological experiments, including analysis and plotting of the data. JvL performed the *ex vivo* recordings, including analysis and plotting of the data. APC performed histological and behavioral experiments. KM and MC conceptualized the project, KM designed the transgenic rat. MC supervised the project and constructed the figures. HB and MC wrote the manuscript. All authors discussed and commented on the manuscript.

## Acknowledgement

We thank Emil Wärnberg for assistance with DeepLabCut. We also thank Pierre Le Merre, Felix Jung, and Katharina Heining for valuable comments on the manuscript. Funding for the study was provided to M.C. by the Wallenberg Scholar program (KAW 2019.0444, Knut and Alice Wallenberg Foundation) and by the Swedish Research Council (**2021-02662).**

## Materials availability

The PV-Cre rat generated in this study is available upon reasonable request to the lead contact.

## Data and code availability

All data and code are available upon request to the lead contact.

## Methods

### Anatomy

All anatomical measurements and nomenclature are based on Paxinos G and Watson C, The rat brain in stereotaxic coordinates, 6^th^ edition. In the current manuscript, mPFC refers to PrL + IL, Cg to Cg1 + Cg2, and dCA1 denotes dorsal CA1 and was determined visually by imaging followed by microscopy.

### Animals

All procedures were approved and performed in accordance and compliance with the guidelines of the Stockholm Municipal Committee. Adult (2-6 months old) male and female heterozygote PV-Cre rats were used. For animals in behavior, the daily amount of food was restricted to 10 g. The animals were weighed daily, and the weight did not subside 85 % of the initial free feeding weight.

### Generation of the PV-Cre rat line

The PV-Cre rat line (‘Pvalb-P2A-Cre rat model’) was produced by SAGE Labs (Saint Louis, MO, USA) using CRISPR-based technology. Cre was introduced into the 3’ UTR of the Parvalbumin (Pvalb) locus using a 2A self-cleaving peptide sequence between Pvalb and Cre. This strategy was chosen to allow the expression of Cre without disruption of the expression of Pvalb.

### Virus injections

Burprenorphine was delivered (I.P 0.05 mg/kg) 30 min before surgery. The animals were deeply anesthetized with isoflurane (1,5-2,5 %), and the body temperature maintained at 37°C using a heating pad. The animals were fixed in a stereotaxic frame (Harvard Apparatus, Holliston, MA) and a small craniotomy (0.5 mm) was made unilaterally over the mPFC, (AP 3-3.3 mm, ML 0.5-0.8 mm). The virus (AAV5-Ef1a-DIO-hChR2(H134R)-mCherry, UNC Gene Therapy Center, #AV4314B, 4x10^12^ virus molecules/ml) was delivered by a glass capillary attached to a motorized Quintessential Stereotaxic Injector (Stoelting, Wood Dale, IL, USA) at rate of 0.05-0.1 μl min^-1^. Injection in the mPFC: half the volume (0.5 μl) was injected at DV -2.0 mm. The capillary was thereafter held in place for 5 min before being slowly lowered to DV –3.0 mm, at which the rest of the virus (0.5 μl) was injected. The pipette was thereafter held in place for 10 min before being slowly retracted from the brain. The incision was closed with stitches (Ethicon, USA), and postoperative caprofren (5 mg/kg) was administered every 24h for 48h.

### Tissue processing

#### General procedure

For perfusions, the animals were deeply anesthetized with pentobarbital and transcardially perfused with 1x PBS followed by 4% paraformaldehyde (PFA) in 1x PBS. The perfused brain was removed from the skull and postfixed in 4% paraformaldehyde in 1x PBS at 4°C for 16 h. The brains were thoroughly washed in 1x PBS and thereafter sectioned (50 μm thickness) using a vibratome (Leica VT1000, Leica Microsystems, Nussloch GmbH, Germany).

#### Immunohistochemistry

PV and Cre co-expression across brain regions and the cortical depth:

Vibratome cut (free floating) tissue was heated to 70°C in 1x Daco Target Retrieval solution (DakoTarget Retrieval Solution, pH 6.1, Dako, code. S2367) for 15 min and left to cool at room temperature (RT). The sections were transferred to 1x TBST (0.3 % Triton-X in 1x TBS) for 1 h, blocked with 10% normal donkey serum in 1x TBST for 1 h, and thereafter incubated with primary antibodies (1:250 Cre-recombinase, rabbit, Synaptic Systems, cat. no. 257 003, and 1:1000 Parvalbumin, guinea pig, Swant, code no. GP72, or Parvalbumin, guinea pig, Synaptic Systems, cat. no. 195 004) in 1x TBST at RT for 12-24 h. The sections were thereafter washed three times in 1x TBST and incubated with a species-specific fluorophore conjugated secondary antibody in 1x TBST for 3-5 h (1:500 donkey anti-rabbit, DyLight 488, Jackson, 711-545-152 and 1:500 donkey anti-guinea pig, Cy3 or Cy 5, Jackson, 706-165-148/706-175-148). The sections were thereafter transferred to 1x TBST with 1:50000 DAPI for 3 min, and consecutively washed with 1x TBST, 1x TBS and 1x PBS (10 min each). The sections were mounted on glass slides (Superfrost Plus, Thermo Scientific^TM^) and coverslipped (Thermo Scientific^TM^) using 50:50 Glycerol:1x PBS.

PV and ChR2-mCherry co-expression in the mPFC:

Immunohistochemistry and mounting of sections were performed as described above. The same PV antibodies were used. Primary antibody for detection of ChR2-mCherry: RFP: 1:500 RFP, rabbit, Rockland code 600-401-379.

### Imaging, and quantifications

For scoring of the co-expression of Cre and PV across brain regions (mPFC, Cg, V1, dCA1, CB; N = 3 PV-Cre rats, CP, GPe; N = 2 PV-Cre rats) and also for the scoring of the co-expression of Cre and PV across the cortical layers (mPFC, Cg, V1; N = 3 PV-Cre rats) tiled z-stack images were acquired at 20x (3-4 sections per brain region for each rat), using a Zeiss (LSM800) confocal microscope with a motorized stage. The co-expression of Cre and PV was quantified using a custom script in R. The position of labelled neurons across cortical layers was determined using a custom MatLab (MathWorks) script utilizing cell and brain surface coordinates manually mapped in ImageJ (Cell counter). Representative high-resolution z-stack images (20-30 focal planes) of neurons co-expressing PV and Cre were acquired at 40x or 63x with the same Zeiss (LSM800) microscope.

Scoring of co-expression of PV and ChR2-mCherrry was done by microscopy of the mounted sections (3-4 sections/rat, N = 3 PV-Cre rats) using a Leica DM6000B fluorescent microscope.

### Patch clamp recordings

For characterization of the neurophysiological properties of cortical neurons expressing Cre and PV, an AAV with Cre-dependent expression (AAV5-Ef1a-DIO-hChR2(H134R)-mCherry (UNC Gene Therapy Center, #AV4314B) was targeted to the mPFC of adult (2-3 months) heterozygote PV-Cre rats (N = 8; 1 μl unilateral injection, 4x10^12^ virus molecules/ml, for details see *Viral injections*). 2-3 w after viral injection, 250 μm thick coronal slices were vibratome cut (VT1200S, Leica, Wetzlar, Germany) in ice cold sucrose solution, containing (in mM): 126 NaCl, 2.5 KCl, 1.25 NaH_2_PO_4_, 24 NaHCO, 25 glucose, 75 sucrose, 1 L-ascorbic acid, 3 Na-pyruvate, 1 CaCl_2_, 4 MgCl_2_. The slices were placed in an extracellular solution, containing (in mM): 126 NaCl, 2.5 KCl, 1.25 NaH_2_PO_4_, 26 NaHCO_3_, 20 glucose, 1.5 CaCl_2_, 1.5 MgCl_2_ and incubated at 34 °C for 30 min, and thereafter kept at RT. For recording, the slices were superfused with extracellular solution kept at 33-35°C. Neurons were visualized using a DIC microscope (Scientifica, Uckfield, UK) with a 60x objective (Olympus, Tokyo, Japan), and fluorescent neurons identified using a pE-400 LED light source (CoolLED, Andover, UK). Patch pipettes (resistance 5-10 MΩ, pulled on a P-87 Flaming/Brown micropipette puller, Sutter Instruments, Novato, CA, USA) were filled with low chloride internal solution containing (in mM): 130 K-gluconate, 5 Na_2_-posphocreatine, 1.5 MgCl_2_, 10 HEPES, 5 Mg-ATP, 0.35 Na-GTP, 1 EGTA, 8 biocytin. Signals were recorded with an Axon MultiCalmp 700B amplifier and digitized at 20 kHz with an Axon Digidata 1550B digitizer (Molecular Devices, San Jose, CA, USA). Access resistance and pipette capacitance were compensated. To assess passive and active membrane properties neurons recorded in current clamp mode were held at a membrane potential of -70 mV. Near-threshold current steps were applied to determine the rheobase current, then 1 s current steps proportional to the rheobase current were applied. An additional small hyperpolarizing step (30 pA) with a duration of 300 ms was used to determine the time constant. To assess the presence of ChR2, a 2 ms blue light stimulation of ChR2 was performed at a holding potential of -70 mV. Electrical properties were extracted using a custom written Matlab (MathWorks, Natick, MA, USA) script. The AP characteristics were determined from the first spike that occurred at rheobase current stimulation. Firing rate and adaptation were determined from a current injection of approximately twice the rheobase current. After recording, slices were fixed overnight in 1x PBS containing 4% PFA. Slices were then washed in 1x PBS, followed by three washes in 0.1M PB containing 0.3% Triton-X (PBT). The slices were incubated at 4°C for one day in PBT with 0.002% Streptavidin conjugated to Alexa Fluor 633 (ThermoFisher Scientific, Waltham, MA, USA), then washed two times in PBT and two times in PB. The slices were mounted and imaged on an LSM800 confocal microscope (Zeiss, Oberkochen, Germany) using a 20x objective.

### Behavioral setup

The setup used for the self-paced reward-seeking task consisted of a linear track (width 8 cm, length 50 cm) with a platform (width, and length 25 cm) at each end. The track and platforms held 2 cm high walls, and the setup was situated 1.5 m above the floor. Two infrared beam break sensors (Adafruit, Product ID: 2168) controlled by an Arduino were used, one tracking entrance into the trigger zone (red time stamp, **Figure 3A**), and one tracking entrance of the rat’s snout into the reward port (red time stamp, **Figure 3A**). All other time stamps were based on video analysis. A spout delivered liquid reward (50 µl 10% sucrose reward, delivered during 2 s) in the reward port situated in the back wall of the reward zone. Reward delivery was triggered by the beam break in the reward port if a beam break had been registered in the trigger zone after the latest beam break in the reward port (= correct trial). In 15% of the correct trials reward delivery was omitted to address trial outcome history.

To accurately align neural activity and behavior, a second Arduino would send pulses (20 Hz) to trigger a Blackfly video camera (FLIR) and to the Digital Lynx 4SX acquisition system, thereby providing a timestamp for each video frame in the electrophysiological recording.

### Behavioral tracking

DeepLabCut (DLC)^57^ was used for tracking animal position. The base of the tail, the end of the tail, and the left and right ears were manually labelled in frames, sampled from all sessions. The behavioral and movement analysis were based on three of the markers tracked by DLC: the base of the tail, and the center of the head (point between left and right ears). Smoothing of tracking data was accomplished using a two-pass Kalman Filter algorithm. The initial state and model parameters were first estimated using an Expectation-Maximization algorithm. A second pass was then performed with a modified observation covariance matrix (multiplied by 5) to reduce the trust in the raw observations, thereby generating smoother estimates.

### Behavioral task

Definition of a trial: leaving the reward zone (= trial start; black time stamp Figure 3A), traversal of the linear track to the trigger zone (traversal ‘to trigger’), turning 180° in the trigger zone, traversal of the linear track back to the reward zone (traversal ‘to reward’), reward consumption/omission, turning 180° in the reward zone, leaving the reward zone (= trial end, black time stamp **Figure 3A**). Reward was randomly delivered in 85% of the trials with successful visit to the trigger zone.

### Behavioral training, including the temporal relation to virus injection, microdrive implantation, and electrophysiological recordings

Behavioral training started 1 w after virus injection. The daily amount of food was restricted to 10 g from the first training day. Animals were weighed daily, and the weight did not go below 85% of the initial free-feeding weight. The first training session the rats could explore the setup and collect reward in the reward port at any point, i.e., no specific behavior was required for reward delivery. When the rats had learned where to collect rewards (>100 collected rewards in 30 mins; usually within the first training session), task training started: the rat was put in the setup and learned through exploratory behavior that visit in the trigger zone was required for reward to be delivered in the reward port. No instructive signals were given to guide the behavior. The rats were trained in one session per day, and rats conducting ≥ 50 trials in 1 h for three consecutive days were considered fully trained and were taken off food restriction. After ≥ three days with free access to food, the microdrive was implanted. After 1 w of recovery from the implantation, the rats were put on food restriction again and three days later task retraining started, with the recording cables attached to the microdrive. The recording sessions started the day after the animal successfully performed 50 trials in 1 h (requiring 1-3 days of retraining) and continued daily until the electrophysiological signal was deemed suboptimal. One day = one session behavior/recording (max 1 h) across the study.

### Microdrive construction

Microdrives (flexDrive^58^, N = 3, or DMCdrive^59^, N = 4) were used for chronic *in vivo* electrophysiology with concurrent optogenetics in behaving adult (3-4 months at experiment start) PV-Cre rats (N = 7 included in analysis). Tetrodes consisted of four twisted fine wires (polyimide insulated ni-chrome wire, 12 μm, Sandvik-Kanthal) that were goldplated to reduce the impedance to 0.2-0.4 MΩ at 1 kHz. Four movable tetrodes were loaded into medical-grade polyimide carrier tubes (0.005 inch OD, Phelps Dodge) in the microdrive. The microdrives were equipped with a Ø200 μm multimode optical fiber (0.5 NA, Thorlabs (FP200ERT)).

### Microdrive implantation

Burprenorphine was delivered (I.P 0.05 mg/kg) 30 min before surgery. The animals were deeply anesthetized with isoflurane (1.5-2.5 %), and the body temperature maintained at 37°C using a heating pad. The animals were fixed in a stereotaxic frame (Harvard Apparatus, Holliston, MA) and a small craniotomy (1.0–1.5 mm diameter) was made unilaterally over the mPFC, (AP 3.0-3.3 mm, ML 0.5-0.8 mm). The microdrive was positioned above the craniotomy with the optical fiber and tetrodes gradually lowered to the PrL (2.5 mm ventral to brain surface), aimed at layer 5/6. Five miniature anchoring screws were used to attach the microdrive to the skull (one contralateral to the microdrive, and two rows of two screws on the anterior and posterior part of the parietal bone). One Teflon coated stainless steel wire (0.005 inch bare, A-M systems) from the electrode interface board (EIB) of the microdrive were connected to the screws for grounding. The microdrive was secured onto the skull using dental adhesive cement (Super Bond C&B, Sun Medical). 3 rats were implanted in the right hemisphere and 4 rats in the left hemisphere. After surgery the animals were single housed. Postoperative caprofren (5 mg/kg) was administered every 24 h for 48 h.

### Neuronal electrophysiological recordings with optogenetics during freely moving behavior

The neural activity was recorded in a total of 97 sessions (N = 7 PV-Cre rats) using a Digital Lynx 4SX acquisition system and Cheetah data acquisition software. The tetrodes were lowered 40-90 μm after every recording session. Unit signals were amplified with the gain of 10,000, filtered with bandwidth 600-6,000 Hz, digitized at 32 kHz, and stored on a PC. LFPs were acquired from one electrode of each tetrode at a sampling rate of 32 kHz. The signal was band-pass filtered between 0.1 and 500 Hz. Single unit and LFP signals were recorded using Cheetah data acquisition system (Digital Lynx 4SX, Neuralynx) in awake animals. Laser light (473 nm, 1-3 ms pulse width, 1s duration, 10 stimulations/frequency, 5-10 mW, 10, 20 and 40 Hz) was delivered through the optical fiber from a DPSS blue laser (Cobolt MLD^TM^ 473 nm, Cobolt) controlled by custom software written in LabVIEW.

### Quantification and statistical analysiS

#### *Ex vivo* electrophysiology

The electrophysiological characterization of PV neurons *ex-vivo* was generated using custom MatLab (MathWorks) scripts.

#### *In vivo* electrophysiology

All analysis of behavior, *in vivo* electrophysiology, and optogenetics was done in python, unless stated otherwise.

#### Cell sorting

Single neurons were manually sorted using MClust (written by A.D. Redish). Well-separated neurons were defined by isolation Distance > 15, L-ratio < 0.2, and the spikes < 0.01 % with < 2 ms^60^.

#### Identification of directly light-activated neurons

To identify directly light-activated FS-PV neurons we employed an optical-tagging test (Stimulus Associated spike Latency Test, SALT)^29^. SALT is used to statistically determine if neurons displaying short-latency spikes with low jitter have significantly changed spike timing in relation to the onset of light pulses. To ensure that the spike sorting was not compromised by light application, the Pearson’s correlation coefficient was used for comparison of waveforms of light-evoked spikes to spontaneous spikes for individual neurons. Using these measures (SALT P-value < 0.05; Pearson’s correlation coefficient r > 0.95) 13 units were identified as FS-PV neurons.

#### Cell classification

The units were first classified into wide-spiking (WS) putative pyramidal neurons and narrow-spiking (NS) putative interneurons based on the distribution of (1) the peak-to-valley ratio (the ratio between the amplitude of the initial peak and the following trough), and (2) the spike-width (time from peak to trough) of each spike waveform, chosen from the tetrode with largest spike amplitude. For objective classification of units, a Gaussian mixture model (GMM) was fitted to the units^61^. Units with NS probability >0.95 were classified as NS, and units with NS probability <0.30 as WS. Units not fulfilling either of these criteria were not classified.

*The waveform similarity index (r):* the pearson’s correlation between the mean spike waveform of each unit (WS, NS, or FS-PV) and the mean light-activated spike waveform.

#### Spectral analysis of the LFP in response to optogenetic stimulation

LFP signals were downsampled to 1KHz and bandpass filtered 1-500 Hz. Relative power was calculated for three frequency bands (6-14, 16-24, and 36-44 Hz) during epochs of light stimulation (1 s) at different frequencies (10, 20 and 40 Hz), and during baseline (1 s preceding light stimulation), respectively. Power spectral density was estimated in frequency range 2-100 Hz using superlets^62^. The relative power was thereafter calculated by dividing the power within each frequency band, to the total power in the power spectrum. The relative power ratio is *P*_light_/*P*_baseline_ where *P*_light_ is the relative power in a frequency band during light stimulation (± 4 Hz around the stimulation frequency) and *P*_baseline_ is the relative power in that band in the absence of light stimulation^35^. The harmonics preclude analysis of how higher frequency bands are affected by lower frequency stimulations is precluded.

#### Spectral analysis of the LFP during running behavior

Raw LFP signals (low-pass filtered at 500 Hz, 32 kHz sampling rate) were processed and analysed using custom software written in Matlab. The LFP signals were downsampled to 1 kHz and LFP power spectra were estimated by the Chronux function (‘mtspectrumc’). To investigate the relative power of the LFP during traversals on the linear track, trials with a more homogenous duration (0.5-1.5 s) of the track traversal (bin 2-9 in Figure 3A) were selected. The power within the frequency band of interest was divided by the total power of frequencies in the 1-100 Hz range and averaged across trials. To analyze the relationship between LFP power and the track traversal or running speed (15-75 cm/sec), LFP power spectrograms (10-100 Hz) were constructed using the continuous wavelet transform (cwt) function with the analytical Morlet wavelet (‘amor’) in Matlab and normalized by the peak power. To identify gamma-band bursts, the LFP power was averaged across the gamma band (30-50 Hz) and then gamma bursts were detected by a threshold (75^th^ percentile power) crossing method using ‘findpeaks’ function in Matlab.

#### Phase coupling analysis

Phase coupling of neuron activity to gamma (30-50 Hz) oscillations was studied for reward and trigger traversals separately. For every spike occurring, the corresponding instantaneous phase of the gamma oscillation was extracted from the Hilbert transform. Only neurons with ≥ 50 spikes in each traversal were included (WS: n = 204, NS: n = 21, FS-PV: n = 13). The strength of phase-coupling of single neurons to gamma oscillations were quantified using pairwise phase consistency^52^ (PPC) (**Figure 7A**) and mean resultant length (MRL; pycircstat. resultant_vector_length) (**Figure 7B**). Significant gamma phase-coupling was based on Rayleigh test (p<0.05, pycircstat.rayleigh).

#### Analysis of single neuron data

Spike data was binned by the recorded frames (50 ms bins; see **Behavioral setup)** and smoothed with a Gaussian filter of 100 ms (scipy.ndimage.gaussian_filter1d) for all analysis described below.

### Generalized linear models (GLMs)

To assess the relationship between mPFC neuronal activity and various behavioral and task-related variables, we employed a Generalized Linear Model (GLM) using a Poisson regressor (sklearn PoissonRegressor). Variables included in the models: categorical variables; *linear position* (the rat’s 1D position in the behavioral setup, in 1 of 10 bins (**Figure 3A**); *run phase* (track traversal to reward zone or trigger zone, respectively), *trial outcome* (correct trial rewarded or not rewarded, respectively), *speed*, *acceleration*, *and turning*, respectively. Additional variables were analyzed but determined to not be significantly encoded: *trial time* (fast vs slow, with cutoff at 3 median absolute deviation (MAD) above median trial time), *trial history* (previous trial rewarded or not), and *trial type* (current trial will be rewarded or not). Run phase was scored specifically during running, i.e., does not include periods of immobility, or turning on a platform **(Figure S3A**). The rat’s turning in the platforms was scored by calculation of the head angular velocity, i.e., the angular difference between the center of the rat’s head and the tail base. The continuous variables speed, acceleration, and turning were z-scored.

In the GLM model the activity of each neuron was represented as a linear combination of the behavioral variables, each scaled by its respective coefficient. To identify which neurons to include in the models, we first fitted the activity of the neurons using models with single variables (**Figure S3B**). Next, we fitted neuronal activity using models with increasing number of variables, until all variables were included. For each model, we quantified the cvR^2^ from the test data (average of held-out data from 5-fold cross-validation) and the training data (average of used data from 5-fold cross-validation), respectively. The predicted explained variance, was computed as the cumulative sum of cvR^2^ from single variable models (**Figures S3B, C)**.

A separate GLM was created for investigation of the relationship between encoding of spatial context and movement variables (speed, acceleration; **Figure 5**). For this an additional variable was generated – position with direction – where position contained 10 bins, with 2 variables for each bin: traversal to reward, and traversal to trigger, yielding 20 predictors, each representing a unique combination of linear position and run phase. The GLMs were evaluated using 5-fold cross validation (Sklearn.model_selection KFold) and the goodness of fit of the prediction was quantified as the squared correlation (Pearson’s correlation (*r*)) between the predicted and the actual test data (sklearn.model_selection cross_val_score).

Two methods were used to quantify encoding of behavioral variables ^38,39^. (I) We created models consisting of single variables and computed the cross-validated explained variance (cvR^2^). This method can overestimate the encoding of a certain variable if that variable is correlated with other variables. Thus, as a second approach (II) we created models with all behavioral variables (full model), followed by shifting one variable in time (circular shift) to generate reduced models. The decrease in explained variance (Δr^2^) for the reduced models, as compared to the full models, reflects the unique encoding of the shifted variable (i.e., the encoding is not captured by other variables). These two measures, thus, provide upper and lower boundaries, respectively, on a given behavioral variable’s explanatory power ^38^.

To examine if the activity of individual neurons correlated to a particular variable, we created reduced models, by shifting one variable in time (circular shift) and quantified the cvR^2^ as described above. To calculate whether the reduction in cvR^2^ was significant, we computed the cvR^2^ for 1000 reduced models. If the cvR^2^ for the full model was greater than 99% of the cvR^2^ derived from reduced models, the neuron was defined as being significantly correlated with that variable.

### Correlations between variable encodings

In figure 3F we correlated (Pearson’s correlation) the unique explained variance (Δr^2^) for all combinations of variables. As a control, Δr^2^-values for each neuron were randomly shuffled prior to computing the correlation between the Δr^2^-values. This step was repeated 1000 times and p-values were computed, by comparing the true correlations with the shuffled control.

### Analysis of spatial activity patterns

To analyze neuronal activity patterns associated with the respective traversal direction, we computed activity vectors as a function of linear position (10 bins, same bins as in GLM). This was done separately for traversals toward the reward zone (run to reward), and from track traversals towards the trigger zone (run to trigger) (**Figure 4**). Traversals longer than 10 s were excluded. Sessions with less than 10 to-reward and 10 to-trigger traversals were excluded, yielding inclusion of 319 neurons (WS: n = 268, NS: n = 38, FS-PV: n = 13). To identify neurons displaying differential activity patterns dependent on the traversal direction, the ‘traversal to trigger’ activity vector was flipped to match the ‘traversal to reward’ activity vector with respect to the start and the end of the traversal. The difference in activity was calculated for each relative position. As control dataset, the traversal direction was randomly shuffled and the difference between the two activity vectors computed 1000 times^63^. Real activity difference that exceeded 97.5% or fell below 2.5% of the shuffled activity differences for was defined as significantly more or less active, respectively.

To compare the mean activity differences for each neuron, based on traversal direction, we computed the direction modulation index (DMI). 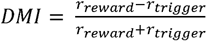, where *r_reward_* is the mean rate of the to-reward activity vector and *r_trigger_* is the mean rate of the to-trigger activity vector. DMI ranges from -1 to 1. Positive values: enhanced spiking in traversals to reward, and negative values: enhanced spiking in traversals to trigger. The value is zero if there is no modulation. Neurons with significant modulation were defined by comparing the trial average mean activity in traversals to reward vs traversals to trigger using Wilcoxon rank-sum (scipy.stats.ranksums).

### Spatial consistency

Spatial consistency was computed by generating activity vectors (see above) for odd and even trials, respectively, followed by correlation of the activity vectors for the odd and even trials (Pearson’s correlation). Control data was generated by shifting (circular shift) the neuronal activities prior to computing the activity vectors, and repeated 1000 times.

### Decoding analysis

We decoded the traversal direction of the rat based on neuronal activity (**Figures 4G,H)**. For each neuron, we computed activity vectors for each track traversal, together with a separate binary vector, describing the traversal direction (0 = to reward, 1 = to trigger).

Using the traversal direction as the label and the vector of activity as corresponding features, we trained support vector machines (SVMs) with linear kernels. For each neuron, we randomly divided 80% of the traversal into a training set and the remaining 20% into a test set. We repeated this step 20 times for each neuron and reported the average accuracy of these 20 iterations as real accuracy. For control, we shuffled the labels prior to training the SVMs. This step was repeated 20 times and the average accuracy of those 20 iterations were reported as shuffled accuracy.

We predicted the traversal direction of the rat, based on neuronal activity, using SVMs trained on activity of different neurons (**Figure 4J)**. Specifically, for each neuron (test neuron) we randomly selected different neurons (training neurons) of same type (with increasing numbers: 1-10 neurons), to train the SVMs. From each selected training neuron, we randomly chose 15 to-reward and 15 to-trigger z-scored activity vectors, and used these activity vectors, together with the direction labels, to train SVMs. Next we evaluated the ability of the SVMs to accurately predict the traversal direction of the rat based on the activity of the test neuron. As a control, we randomly shuffled the labels prior to training the SVMs. This procedure was repeated, such that all neurons were used as test neurons.

### Correlation between unique encoding and decoding performance

We correlated the decoding performance with the Δr^2^ (See GLM section) for each behavioral variable in the model (**Figure 4I)**. As control, we generated shuffled data by shuffling the Δr^2^ values for each neuron, prior to computing the correlation between decoding performance and Δr^2^ values. The shuffling was repeated 1000 times.

### Spatial Information

To assess the extent to which the activity of neurons in the PrL is linked to the spatial location of the track that the animal is occupying and the direction of the traversal, we calculated the spatial information (SI) for each neuron for both directions separately (**Figure 6I**). SI was calculated as 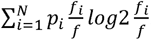, where *N* is the number of bins, *p_i_* is the occupation probability in the *i*^th^ bin, *f_i_* is the mean activity in bin *i*, and *f* is the average activity of the neuron across all bins^5^.

### Statistics

Across all figure panels, the significant differences are indicated for all analyses performed (*p < 0.05, **p <0.01, ***p < 0.001), i.e., non-significant (ns) differences are not indicated.

