## Supplementary information for "Cell type-specific representation of spatial context in the rat prefrontal cortex"

**Figure S1. Additional characterization of the PV-Cre rat line**

(**A**) Example of immunohistochemical detection of Cre (green) in prefrontal PV neurons (red) in an adult PV-Cre rat. Nuclei counterstained with DAPI (cyan).

(**B**) Split channels of the immunohistochemical detection of Cre (green) in PV neurons (red) in the cerebrum and cerebellum of adult PV-Cre rats presented in Figure 1B. Nuclei counterstained with DAPI (cyan).

(**C**) Investigation of the specificity and efficiency, respectively, of targeting of Cre to PV neurons throughout the cortical depth in three different cortical regions (Cg, mPFC, and V1). Gray: specificity, i.e., the fraction Cre-expressing neurons also expressing PV. Red: efficiency, i.e., the fraction PV neurons also expressing Cre.

(**D**) Box plots of additional intrinsic properties of the recombined mPFC PV neurons recorded with whole-cell patch-clamp electrophysiology *ex vivo* (Figures 1D-H). Rheobase current: 206 ± 70 pA, time constant (tau): 4.9 ± 1.5 ms, AP downstroke: 0.44 ± 0.07 ms, AP threshold: -40 ± 3 mV (mean, s.d.). All properties are typical of cortical FS-PV neurons.

Scale bars: 500 μm (A), 20 μm (B); mean ± SEM (C); boxplots (D): black bar: median, box: 25th-75th percentile, whiskers: 99.3% data coverage.

**Figure S2. Electrophysiological recordings in awake PV-Cre rats: optotagging, induction of gamma oscillations, and behavior.**

(**A**) Anatomical outline of the reconstructed recording sites in the PV-Cre rats (N = 7) included in the electrophysiological analyses.

(**B**) Example coronal section from one (133655_3) of the seven PV-Cre rats. Red: ChR2-mCherry expression in mPFC FS-PV neurons. White arrowheads: electrolytic lesions used for reconstruction of the last recording site.

(**C-D**) Detailed analysis of the behavior in the self-paced reward-seeking task during electrophysiological recordings. 1 recording session: 100 trials, or max 1 h. Session time for implanted rats (N = 10): 24 ± 9 minutes/session.

(**C**) Trials per session (n = 78 ± 27), and the total number of trials (numbers in bars) for all implanted PV-Cre rats (N = 10). Four rats conducted a comparably low number of trials per session, and we therefore analyzed the temporal variability of the individual trials (D) to catch suboptimal trial behavior. Dashed line: mean number of trials per session across the animal cohort.

(**D**) Trial time, identifying highly varied trial behavior for three rats (132456_3 (green), 373586_5 (pink), and 132461_2 (red)). These rats in addition did few trials per session and < 500 trials in total (C) and were therefore excluded from electrophysiological analyses.

(**E**) Left: for objective classification of mPFC neurons into WS and NS, a gaussian mixture model (GMM) was fitted to the spike width and the peak-to-valley ratio (PVR) of the individual neurons. This revealed the NS probability (gradient bar) of individual neurons. Right: the classification of WS, and NS, neurons, respectively, based on the GMM clustering. Units with high NS probability (> 0.95) were classified as NS (n = 54; blue) and units with low NS probability (< 0.3) were classified as WS (n = 281; gray). Neurons with intermediate NS probability were not classified (n = 14; green) and discarded from future analysis. All optotagged FS-PV neurons (n = 13; black circles) as expected belonged to the NS population.

(**F**) Example single LFP traces during baseline (-1 to 0 s) and light stimulation (0 to 1 s) at 10 (left), 20 (middle), or 40 Hz (right). Top: raw LFP (0.1-500 Hz). Bottom: spectrogram (2-100 Hz). All three stimulation frequencies evoked a response in the LFP visible as a local maximum at the stimulation frequency and its harmonics.

(**G**) The mean relative LFP power (2-100 Hz) during baseline and in response to light stimulation at 10 (left), 20 (middle), or 40 Hz (right). All stimulation frequencies increased the LFP power around the stimulated frequency and its harmonics. For number of recordings, see (H).

(**H**) The relative LFP power ratio in response to 10, 20, and 40 Hz activation of mPFC FS-PV neurons for all individual recording sessions.

Scale bars: 500 μm (B); mean ± SEM (C, D, G).

**Figure S3. Variable selection for GLMs**

(**A**) Example scoring of the running behavior of one rat during 30s. During turning and reward consumption in the reward zone (part of bin 10) and during turning in the trigger zone (part of bin 1), the behavior is scored as ‘Not running’. The direction of the running (to reward vs to trigger) defines the variable *run phase.*

(**B-C**) Selection of behavioral and task variables for the GLMs.

(**B**) Explained variance (mean across neurons, n = 322), sorted in descending order, of linear models containing single behavioral or task variables.

(**C**) Explained variance (mean across neurons, n = 322) of linear models containing increasing numbers of variables. Brown: the explained variance of the test data, orange: the explained variance of the training data, gray: the cumulative sum of the explained variance of single variable models, representing the predicted explained variance if all variables are independent. The variables significantly increasing the explained variance of the test data (black horizontal bar) were included in the full model.

(**D**) Calculation of the unique explained variance (∆r^2^) i.e., the difference between the explained variance of the full model (x-axis) and of the reduced models (y-axis). Scatter plots of the mean explained variance of the full models (x-axis) vs the respective reduced models (y-axis) for all individual neurons.

(**E**) The unique explained variance (mean across all neurons, n = 322) of the variables included in the full model.

Mean ± SEM (B, C, E); *p < 0.05, **p < 0.01, ***p < 0.001.

**Figure S4. Consistent spatial representation across the PrL neurons**

(**A**) The trial-averaged FR (z-scored) of all individual neurons at the 10 track positions. The data was split into two groups based on the traversal direction (to reward vs to trigger), and each group was split into two based on odd vs even trial number. Neurons were sorted by the peak FR in odd trials and plotted in the same order in the even trials.

(**B**) The similarity of the FR between odd and even trials (Pearson’s correlation (*r*)) was significantly higher for the real data compared to the shuffled data (circular shifted spike trains) (p = 5.1×10^-138^, paired t-test).

(**C**) The FR in individual trials for one example neuron of each cell type.

***p < 0.001.

**Figure S5. The modulation of PrL gamma oscillations during reward-seeking is not related to speed processing**

(**A**) Top: the mean power (normalized) in the LFP frequency spectrum (10-100 Hz) across the 91 recording sites during traversals to reward (left) and to trigger (right), respectively. Bottom: extraction of the mean power in the gamma band (30-50 Hz). µ = mean traversal time.

(**B**) The difference in LFP power (normalized, 10-100 Hz) across the linear track between traversals to trigger and traversals to reward.

(**C**) Movement trace based on point at center of the head (gray) of one rat during a single session, tracked by DeepLabCut. Orange: detection of the rat’s position on the linear track (bin 2-9 in Figure 3A) during traversal to the reward platform; purple: detection of the rat’s position on the linear track (bin 9-2 in Figure 3A) during traversal to the trigger platform.

(**D**) Example detection of gamma bursts (red dots) in a single LFP trace (0.8 s) band-pass filtered on gamma frequencies (30-50 Hz). Dashed line: the peak detection threshold (75^th^ percentile power).

(**E**) The distribution of speeds detected in the 91 recording sites (speed bin size = 1 cm/s). To adjust for the differential speed in the two track traversals, only speed bins within both traversals’ speed distribution (15-75 cm/s, gray dashed vertical lines) were included in the LFP analysis (Figures 6G, S6F, G). Dotted lines: the mean number of recordings sites included across the speed bins for traversals to reward (orange) and to trigger (purple), respectively.

(**F**) The relationship between the mean LFP power (normalized, 10-100 Hz) and the running speed (15-75 cm/s) in traversals to reward (left) and to trigger (right). Dotted line: the frequency (30-80 Hz) with the maximum power in each speed bin.

(**G**) The difference in the relationship between the LFP power (normalized, 10-100 Hz) and the speed between traversals to trigger and traversals to reward.

91 recordings sites (A, B, E, F). Start/end = bin 2 or 9, depending on the traversal direction (see Figure 3A); mean ± SEM (A, bottom).
