## Supplementary figures and images for "Cell type-specific representation of spatial context in the rat prefrontal cortex"

### Supplementary figures

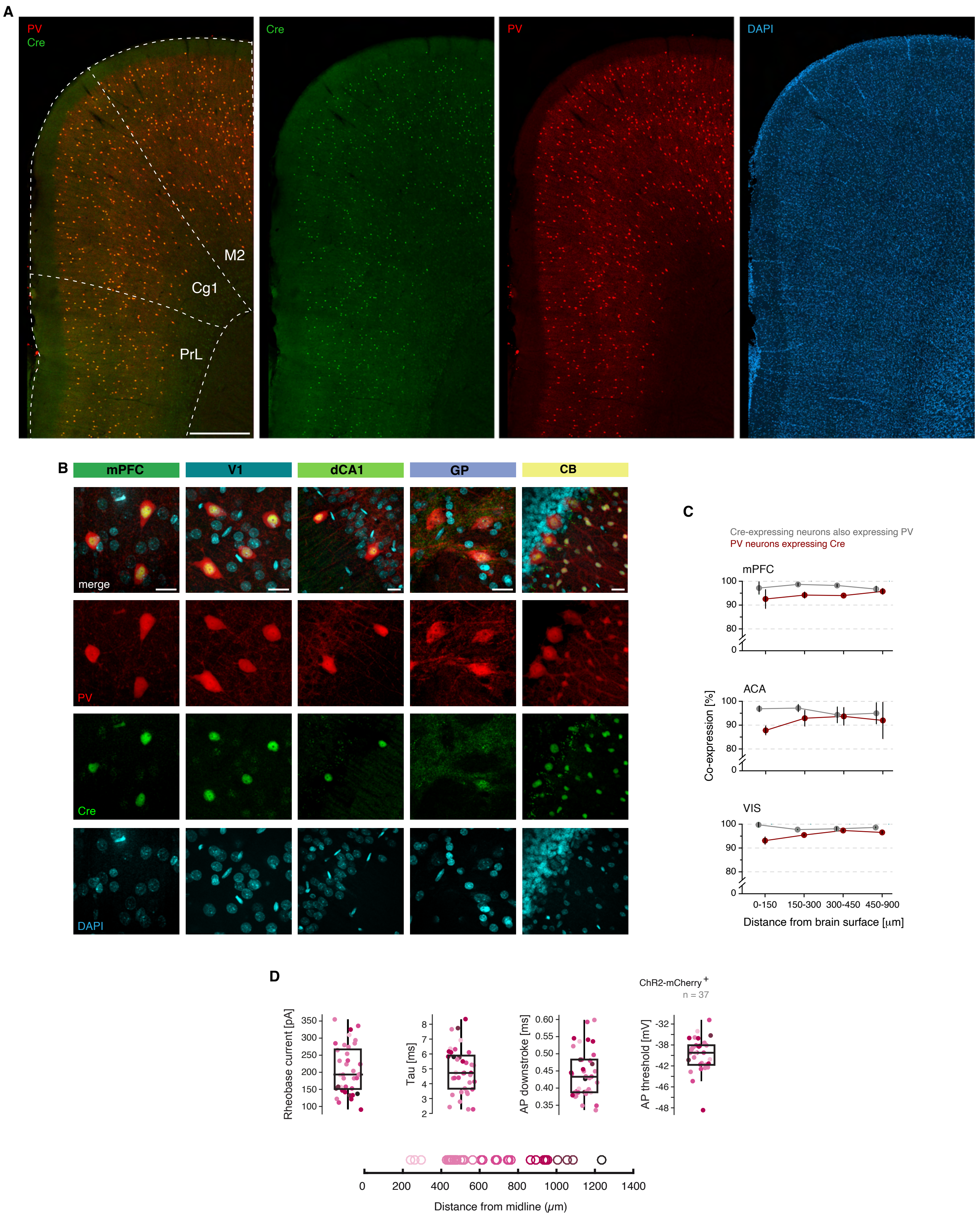

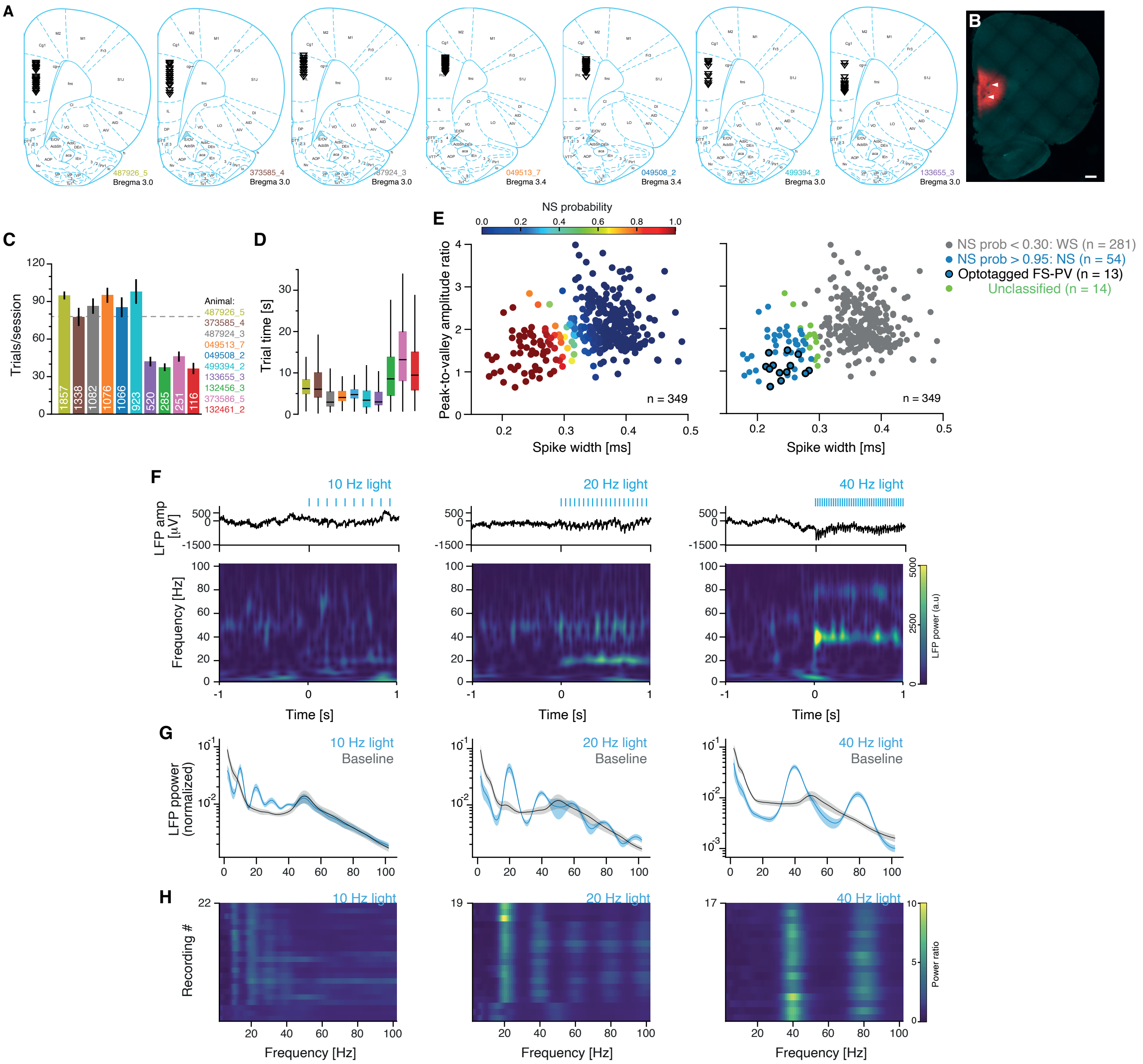

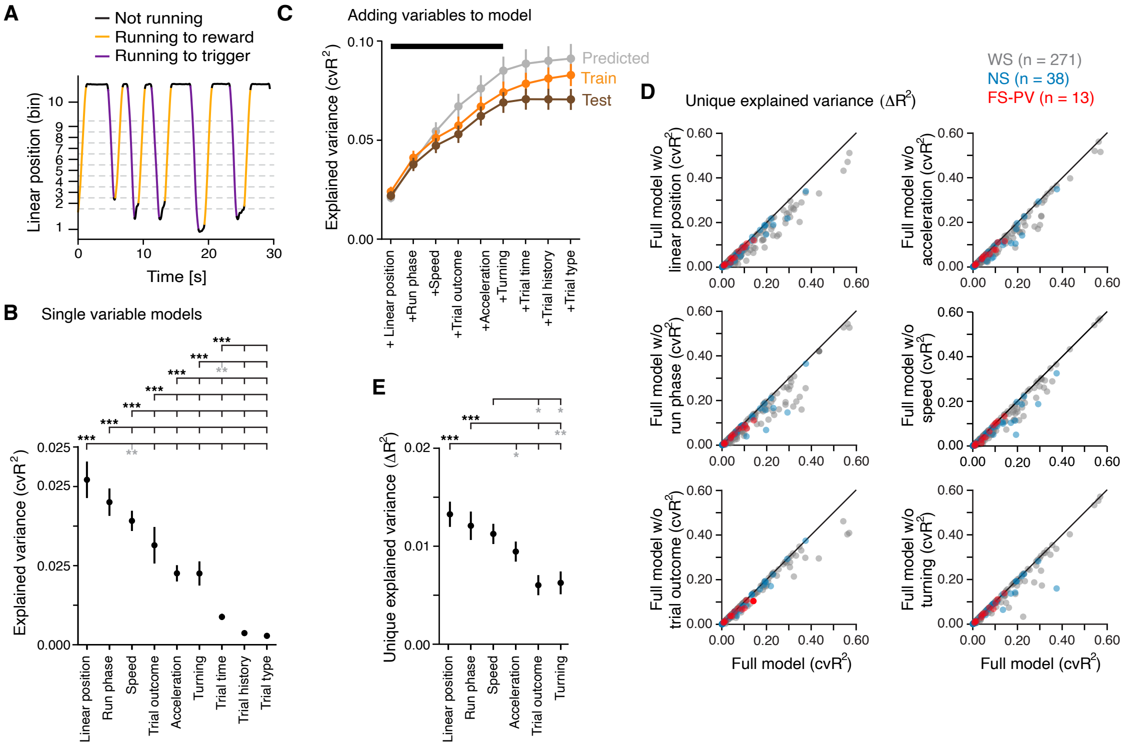

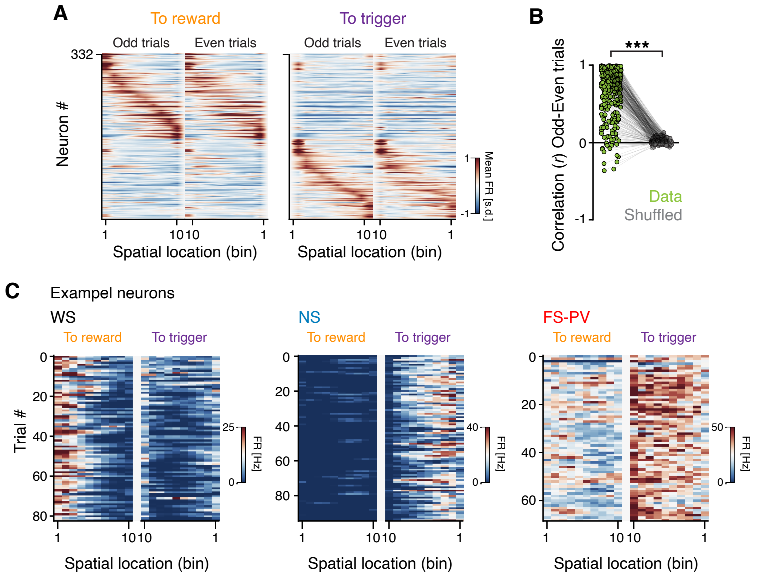

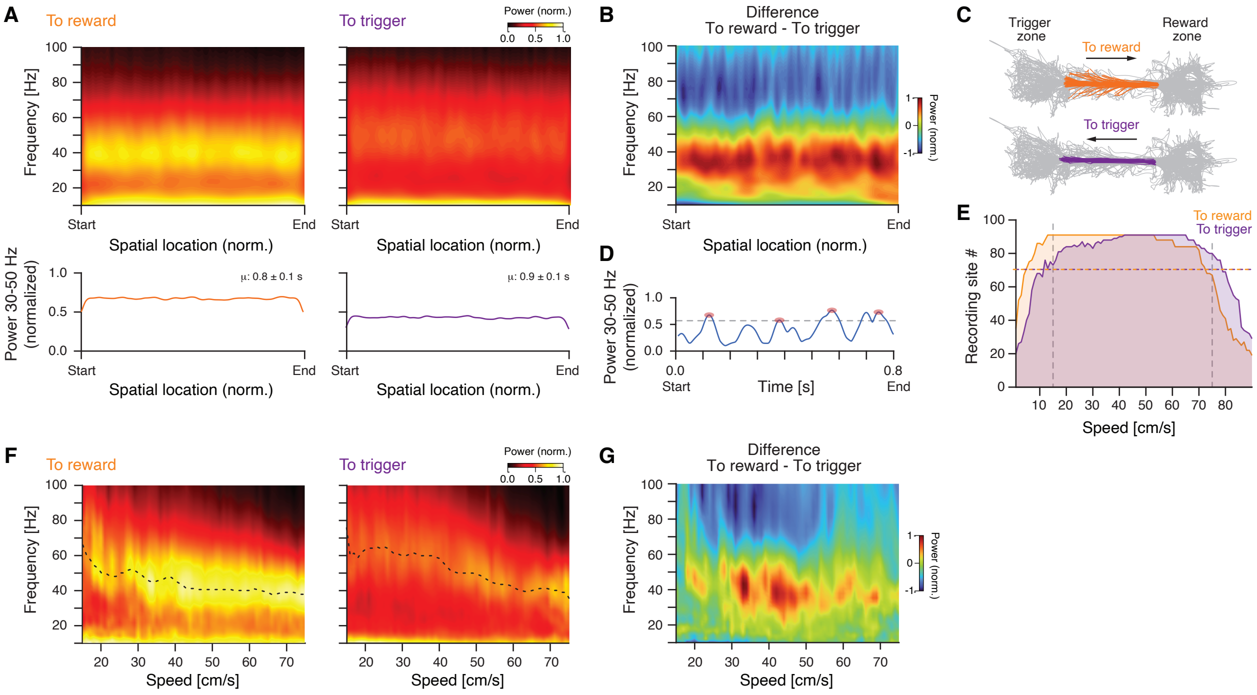
